# Loss of Neurodevelopmental Gene *CASK* Disrupts Neural Connectivity in Human Cortical Excitatory Neurons

**DOI:** 10.1101/2022.02.14.480404

**Authors:** Danny McSweeney, Rafael Gabriel, Kang Jin, Zhiping P. Pang, Bruce Aronow, ChangHui Pak

**Affiliations:** Graduate program in Molecular and Cellular Biology, UMass Amherst, Amherst, MA 01003 USA; Department of Biochemistry and Molecular Biology, UMass Amherst, Amherst, MA 01003 USA; Depts. of Biomedical Informatics, Pediatrics, University of Cincinnati, Cincinnati Children’s Hospital Medical Center, Cincinnati, OH 45229 USA; Child Health Institute of New Jersey and Department of Neuroscience and Cell Biology, Robert Wood Johnson Medical School, Rutgers University, New Brunswick, NJ 08901 USA

**Keywords:** Schizophrenia, synapse, iPS cells, stem cells, neurexin, cell adhesion, synaptic transmission, disease modeling, neuroligin, neurotransmitter release

## Abstract

Loss-of-function (LOF) mutations in *CASK* cause severe developmental phenotypes, including microcephaly with pontine and cerebellar hypoplasia, X-linked intellectual disability, and autism. Unraveling the pathogenesis of *CASK*-related disorders has been challenging due to limited human cellular models to study the dynamic roles of this molecule during neuronal and synapse development. Here, we generated *CASK* knockout (KO) isogenic cell lines from human embryonic stem cells (hESCs) using CRISPR/Cas9 and examined gene expression, morphometrics, and synaptic function of induced neuronal cells during development. While young (immature) *CASK* KO neurons show robust neuronal outgrowth, mature *CASK* KO neurons displayed severe defects in synaptic transmission and synchronized burst activity without compromising neuronal morphology and synapse numbers. In developing human cortical neurons, CASK functioned to promote both structural integrity and establishment of cortical excitatory neuronal networks. These results lay the foundation for future studies to identify suppressors of such phenotypes relevant to human patients.

**Highlights:**

- *CASK* LOF mutations increase neuronal complexity in immature developing neurons
- *CASK* LOF does not alter synapse formation and neurite complexity in mature neurons
- Synaptic transmission and network synchronicity are compromised in *CASK* KO neurons
- Differential gene expression analysis reveals enrichment of synaptic gene networks in mature *CASK* KO neurons

## Introduction

Defining the molecular and cellular mechanisms underlying neurodevelopmental disorders (NDDs) is crucial for developing novel therapeutics and treatment of patients with these devastating conditions. Advances in genome-wide association studies and identification of genetic variants associated with NDDs, even those with heterogeneous polygenic origin, have expanded new fertile grounds for disease mechanistic studies linking genetic variants to human phenotypes. *CASK*-related disorders present such a case. Located on chromosome Xp11.4, *CASK* encodes a multi-domain scaffolding molecule named calcium/calmodulin-dependent serine protein kinase (Hata et al., 1996; Hsueh, 2009). To date, multiple genetic variants of *CASK* have been identified in the human disease population, including microcephaly with pontine and cerebellar hypoplasia (MICPCH), X-linked intellectual disability (XLID), FG syndrome, developmental delays and autism spectrum disorders (ASDs) (Froyen et al., 2007; Najm et al., 2008; Piluso et al., 2003; de Vries et al., 2002). Despite the rich human genetic findings, our understanding of how *CASK* loss-of-function (LOF) mutations impact neurodevelopment and function in the developing human brain is lacking. This challenge stems from the limited access to patient human fetal brain tissue and the pleiotropic and multi-faceted nature of *CASK* function in the nervous system.

CASK is ubiquitously expressed in the brain with the ability to bind to and work with multiple proteins at various cellular compartments over the course of development. In the developing rodent neurons, CASK shuttles into the nucleus to regulate gene expression of N-methyl-D-aspartate receptor (NMDAR) subunit 2b (*NR2b*) and *Reln* by binding to T-box transcription factor Tbr1 and CNIAP (CASK-interacting nucleosome assembly protein) (Hsueh et al., 2000; Wang et al., 2004a, 2004b). During synapse development, CASK’s prominent role at the presynapse is as a scaffolding molecule where it interacts with N-type calcium channels, adapter proteins (Veli, Mint, Liprin-alpha), and cell adhesion molecules (Neurexin-1), thereby maintaining the molecular architecture for neurotransmitter release (Butz et al., 1998; Hata et al., 1996; LaConte et al., 2016; Mukherjee et al., 2008; Olsen et al., 2005; Spangler et al., 2013). At the postsynapse, CASK interacts with cell adhesion molecule Syndecan-2 to regulate the stability/maintenance of dendritic spines via downstream signaling events mediated by NF1-PKA-Ena/VASP pathway (Chao et al., 2008; Lin et al., 2007). CASK also functions as a bridge between the plasma membrane and the F-actin cytoskeleton via binding to protein 4.1 (Chao et al., 2008). Furthermore, in *Drosphila*, CASK physically binds to CaMKII and regulates the phosphorylation status of CaMKII, suggesting that CASK acts as an activity sensor controlling a major signaling hub at the synapse upon neuronal activity (Hodge et al., 2006; Lu et al., 2003).

Despite these illuminating studies linking CASK to cortical development and synapse maturation, animal and human cellular models of *CASK* LOF effects have been elusive. Although the diversity of CASK function has been demonstrated through biochemical studies, many of these interactions have been performed with limited domains and lack the cellular context details, which makes it difficult to pinpoint exactly what its normal function is during brain and synapse development. Moreover, while invertebrate studies strongly support the role of CASK in synaptic transmission at neuromuscular junctions (Chen and Featherstone, 2011; Zordan et al., 2005), *Cask* KO in mice leads to perinatal lethality, and, in surviving animals, subtle changes in synaptic transmission (increased excitatory mini postsynaptic currents and decreased inhibitory postsynaptic currents) with normal ultrastructure of synapses, suggesting that the function of this molecule can be different in vertebrates (Atasoy et al., 2007). More recently, novel genetic mouse models using X-inactivation scheme report defects in excitatory/inhibitory balance and decreased GluN2B mRNA levels in heterozygous *Cask* KO mosaic females (Mori et al., 2019). This defect was primarily rescued by restoring GluN2B, suggesting a postsynaptic mechanism (Mori et al., 2019). In comparison, human iPSC-neurons derived from two different patients carrying *CASK* mutations reveal some deficits in inhibitory presynaptic puncta size in one patient background (Becker et al., 2020). However, due to heterogeneity in patient genetic background, sex, and nature of *CASK* mutations, patient iPSC-derived neuronal studies require multiple lines and proper controls. Therefore, there is a critical need for a study with isogenic lines for *CASK* LOF, which would provide a clean genetic background from which one can assess the effects of *CASK* LOF without the influence of individual’s genetic background.

Here, we describe an approach using CRISPR/Cas9 gene editing in hESCs to generate two independent *CASK* KO isogenic lines. Using wildtype unedited hESC line as a control, we performed molecular- and cellular-based analyses at two different developing time points during neuronal differentiation and synapse formation (day 7 vs. day 28). Consistent with CASK’s predicted role in dendritic morphogenesis, we observed that immature neurons show an increased dendritic complexity and upregulation of gene networks related to neuronal cell adhesion, neurite outgrowth, and cytoskeletal organization. Interestingly, upon maturity and well into synapse development, mature neurons lacking *CASK* showed defects in neuronal spiking and synchronized burst firing patterns without significant changes in neuronal morphology and synapse numbers. Whole-cell patch electrophysiology measurements revealed that *CASK* LOF results in a selective decrease in spontaneous synaptic event frequency without changing intrinsic membrane properties or action potential firing, which suggests a presynaptic defect. In agreement with this result, differential gene expression analysis by bulk RNA sequencing revealed significant changes in gene regulatory networks associated with synaptic function. In summary, while CASK functions to stabilize dendritic outgrowth in developing human cortical neurons, at post-neurogenesis CASK plays an essential role in synaptic release and establishment of cortical excitatory neuronal networks without directly influencing synapse formation. These results emphasize that human neuronal models of *CASK* deficiency can provide a useful cellular tool to elucidate the molecular and cellular mechanisms underlying *CASK*-related syndromes.

## Results

### Generation of isogenic CASK KO hESC lines and differentiation into cortical excitatory induced neurons

Using H1 hESCs as the parental male control line, we introduced a plasmid (lenti-CRISPR v2) expressing Cas9 and *CASK*-specific guide RNAs (sgRNAs) targeting the first coding exon via nucleofection (Fig. 1A). Transient selection by puromycin allowed propagation of hESCs that carried this construct. Upon replating this edited pool, we established single cell clones which were screened by PCR amplification and sanger sequencing (Fig. 1C). Through this method, we established two independent *CASK* null cell lines (KO #1 and KO #2), which represent 14 bp and 10 bp deletions, respectively. These deletions disrupt the first coding exon, resulting in a frameshift and a complete LOF for CASK protein as verified by immunoblotting (Fig. 1D). Both KO #1 and #2 hESCs were able to differentiate efficiently into cortical excitatory neurons by NGN2 forced expression and were chosen for subsequent molecular and cellular analyses.

**Figure 1.**
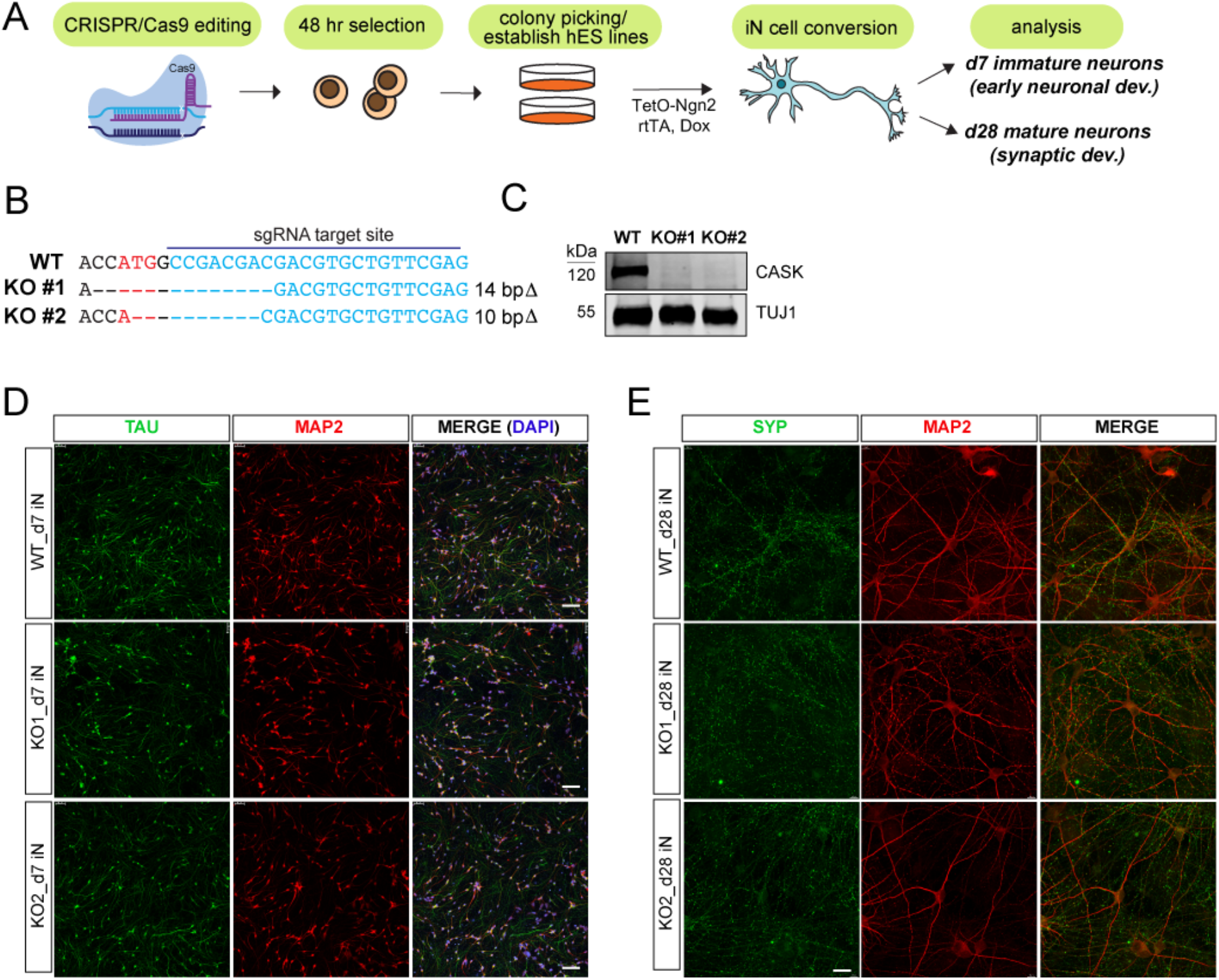
CRISPR-mediated generation of *CASK* KO hESC lines and differentiation into iN cells. (A) A schematic of workflow describing the generation of constitutive *CASK* KO hESC lines via CRISPR/Cas9, expansion of clonal cell lines and Ngn2-iN cell differentiation. Analysis time points were at day 7 to assess early neuronal development and at day 28 to assess synaptic development. (B) Verification of gene edited genomic DNA sequence in the human *CASK* exon 1 by Sanger sequencing. sgRNA target region downstream of ATG site (red) is noted. Resulting deletions obtained using CRISPR-mediated gene editing are shown. (C) Immunoblot verification of complete *CASK* LOF in day 4 Ngn2 iN cells using an anti-CASK antibody. TUJ1 was used as a loading control. (D) Representative confocal images of immunostained d7 immature iNs showing developing axons (TAU-green) and dendrites (MAP2-red) (scale bars, 100 μm). (E) Representative confocal images of d28 iN cells immunostained with a pre-synaptic marker SYP (green) and a dendritic marker MAP2 (red) (scale bars, 20 μm).

### Differential gene expression networks in CASK KO immature neurons

Since CASK has been shown to play roles both as an early neurodevelopmental effector and a late synaptic organizer (Hsueh, 2009), we chose an early development time point (day 7) to investigate the neurodevelopmental effects of *CASK* LOF in human cortical neurons without the presence of supporting mouse glial cells (Zhang et al., 2013). Taking total RNA from each differentiated cell lines, we performed bulk RNA sequencing to examine any potential gene expression program differences between WT and KO #1 and #2 genotypes in early developing neurons. Four independent culture replicates were used for this analysis. Through differential gene expression analysis using DESeq2 (cutoff of FC ≥ 1.2 or FC ≤ 0.8, p ≤ 0.05; minimum of 5 TPM per gene), we found a total of 876 number of DEGs with 420 as upregulated and 456 as downregulated (Fig. 2A, see SI). Unbiased gene set enrichment analysis (GSEA; ToppGene,(Chen et al., 2009)) revealed that a significant number of upregulated DEGs concentrated on cell adhesion (p value 5.85E-06), cell projection and morphogenesis (p value 9.60E-07), neuronal development (p value 1.26E-05), and synaptic membrane/structure (p value 1.56E-08) (Fig. 2B). This pattern of enrichment was far minimal in the downregulated DEGs (see SI).

**Figure 2.**
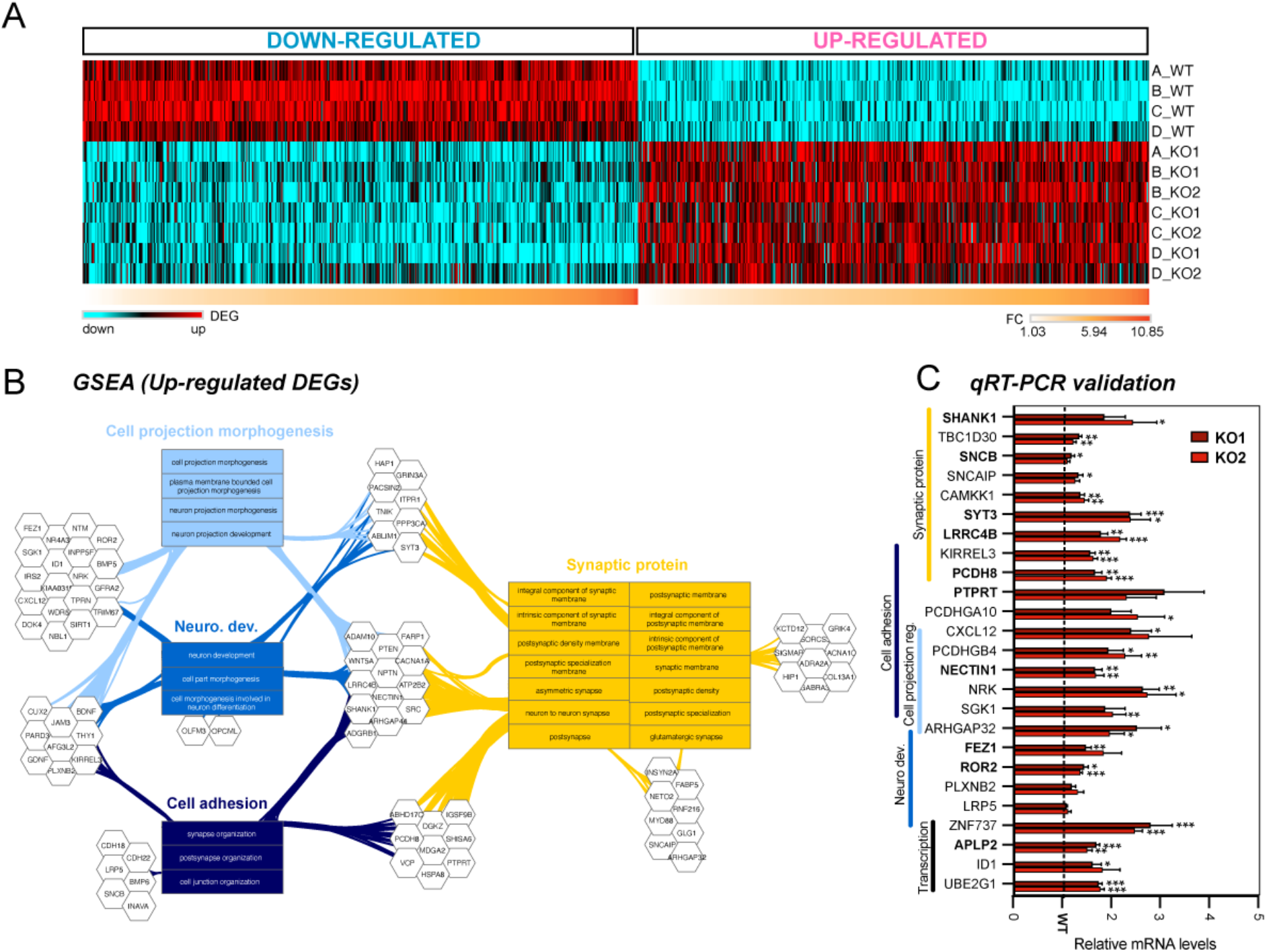
Differential gene expression analysis in d7 *CASK* KO immature iNs using bulk RNA-sequencing. (A) Heatmap showing both up- and down-regulated DEGs across independent culture replicates (4 replicates for WT & KO#1, 3 replicates for KO#2). Cutoff for DEGs was set at FC ≥ 1.2 or <=0.8, p<0.05; minimum of 5 TPM per gene). A total of 876 shared DEGs were identified in CASK KO#1 and KO#2, as compared to WT control, with 420 genes upregulated and 456 genes downregulated. (B) Gene set enrichment analysis (GSEA; ToppCluster) of commonly upregulated DEGs (Bonferroni p<0.05) identified 3 major biological pathways (BP; cell projection morphogenesis p<9.60E-07; neurodevelopment p<1.26E-05; cell adhesion; p<5.85E-06) and 1 cellular component (CC; postsynapse; p<2.01E-08) affected by *CASK* LOF. (C) qRT-PCR validation of specific up-regulated DEGs for both KO#1 and KO#2 relative to WT (set to 1). GAPDH probe was used as a normalization control. Bolded probes indicate genes cross-referenced with SynGO. Four independent replicates per genotype were validated, and each qRT-PCR experiment was performed in triplicate for 26 upregulated DEGs. qRT-PCR data represents means ± SEM, and statistical analysis was performed using Student’s t-test comparing WT to each individual KO (*P<0.05, **P<0.01, **P<0.001; nonsignificant comparisons are not indicated).

In order to validate these results, we performed quantitative RT-PCR using probes against 25 up-regulated DEGs and 12 down-regulated DEGs from total RNA isolated from 4-5 independent culture samples. A vast majority of these hits were validated with statistical significance confirming these genes as target DEGs (Fig. 2C, Fig. S1). Among these targets, multiple synaptic genes (SynGO, (Koopmans et al., 2019)) were validated including *SHANK1, SNCB, SYT3, PCDH8, PTPRT, NECTIN1, FEZ1, ROR2, APLP2, CNTN2, GDI1,* and *RPL10*. In addition, genes related to cell adhesion and structural regulation (*PCDHGA10, CXCL12, NRK, ARHGAP32, PLXNB2*, and *LRP5*) and transcription (*ZNF737, ID1*, and *UBE2G1*) were also validated. These results show that there exists dysregulated neuronal organization in the absence of *CASK*, which highlight its cell autonomous role in controlling early neuronal outgrowth and morphology.

### CASK KO neurons display neurite overgrowth at an early stage of development in culture

Normally, active neuronal growth occurs in day 7 NGN2 neurons as they undergo dendritic and axonal outgrowth and exhibit gene expression profiles of immature, developing neurons (Chanda et al., 2019; Zhang et al., 2013). Compared to WT, day 7 neurons lacking *CASK* showed gene expression profiles indicative of neuronal overgrowth and upregulated complexity, which prompted us to examine their morphology. Using transient transfection at day 4, neurons were sparsely labeled with green fluorescent protein (GFP) expressed from Synapsin promoter (SYN-EGFP vector). At day 7, we fixed these cells and performed confocal imaging and neurite outgrowth analysis to measure various parameters as previously performed (Pak et al., 2015, 2021). Compared to WT, *CASK* KO neurons showed an overall increase in total dendritic length and number of branch points without significant changes in soma size and the number of primary processes (Fig. 3A, B). These results are in agreement with gene expression data in that, in the absence of *CASK*, neurons fail to restrict neuronal complexity and structural integrity in immature human neurons.

**Figure 3.**
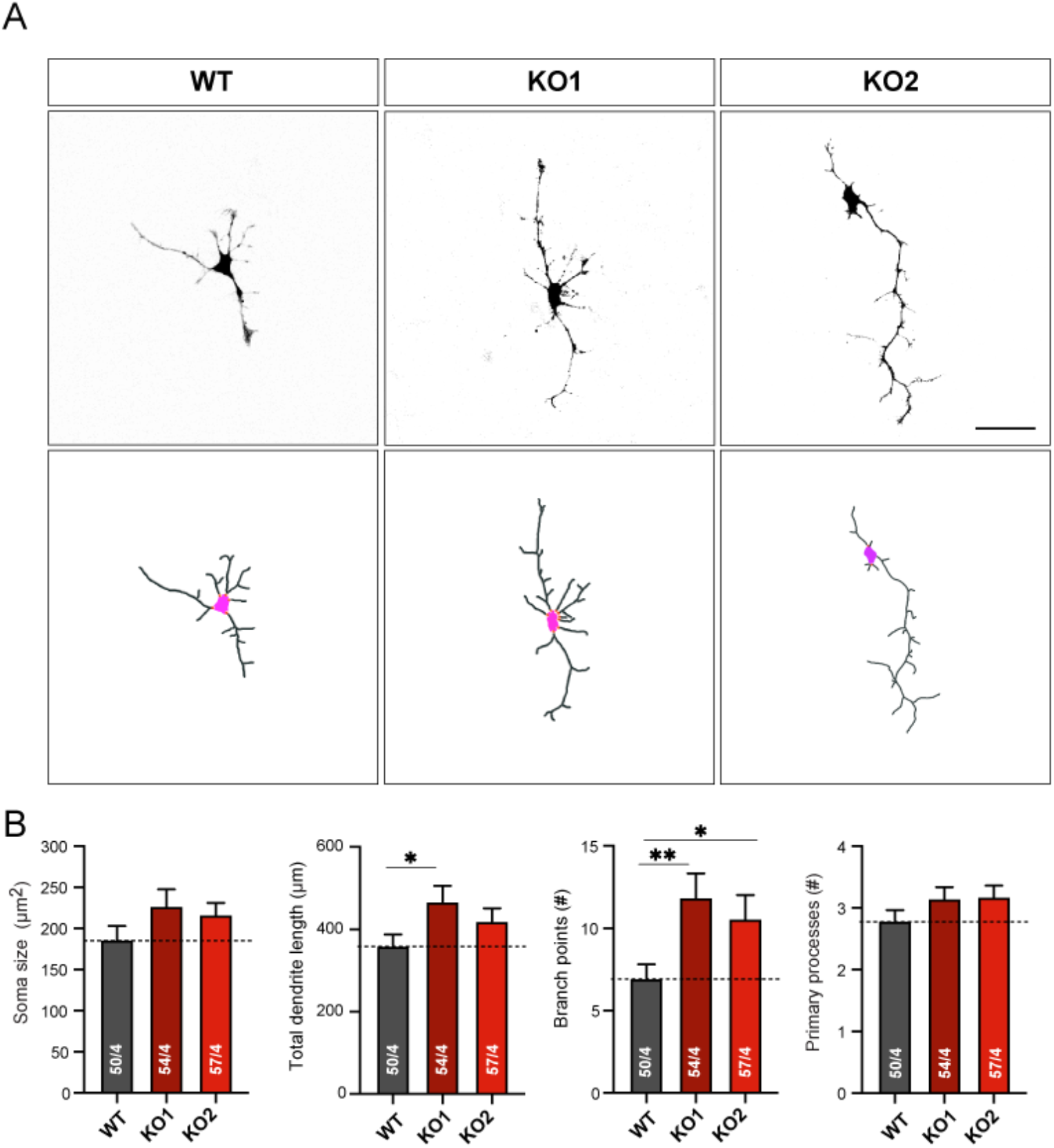
Quantification of neurite outgrowth for immature d7 *CASK* KO Ngn2 iNs shows an overgrowth phenotype. (A) Representative images (top) and binary mask (bottom) of day 7 Ngn2 iNs which were transfected with SYN-EGFP construct at day 4 to label single neurons for morphological quantification (scale bar, 50 μm). (B) Quantification of morphological parameters in WT and KO#1 and KO#2 using Imaris (Bitplane). Summary graphs of soma size, total dendritic length, number of branch points, and number of primary processes. Data represents means ± SEM (numbers in bars represent # of cells/# of independent culture replicates performed). Statistical analysis was performed using Student’s t-test comparing WT to each individual KO (*P<0.05, **P<0.01; nonsignificant comparisons are not indicated).

### Mature neurons lacking CASK exhibit normal neuronal and synaptic morphology

In order to understand the functional role of CASK in synaptic development in human cortical neurons, we co-cultured WT and *CASK* KO day 3 NGN2 neurons with mouse glia and assayed neuronal morphology at day 28. Day 28 is a relatively mature time point with well documented active synaptic properties and elaborate neuronal morphology (Pak et al., 2021; Zhang et al., 2013). Similar to day 7 analysis, we transiently transfected developing neurons with SYN-EGFP plasmid at day 14 to sparsely fill neurons with EGFP and performed immunohistochemistry using antibodies against the presynaptic marker Synaptophysin (SYP) and the postsynaptic marker PSD-95. By confocal imaging, we acquired single neuron images at 20X to quantify neurite outgrowth and focused dendritic segments at 60X to quantify synapse numbers. From four independent culture batches, we quantified various parameters including soma size, total dendritic length, number of branch points, and number of primary processes for neurite analysis and synaptic density and volume for synapse analysis. As shown in Fig. 4, there was no significant change in the parameters measured for neurite complexity except a slight decrease in soma size for KO #1 compared to WT (Fig. 4A, B). Overall, the complexity of *CASK* KO neurons were similar to those of WT neurons, unlike what we have observed in day 7 neurons. In synaptic puncta measurements, we found no difference in the density or the volume of SYP and PSD-95 puncta on fixed dendritic segments, suggesting that synapse formation is not altered in *CASK* KO cells (Fig. 4C, D).

**Figure 4.**
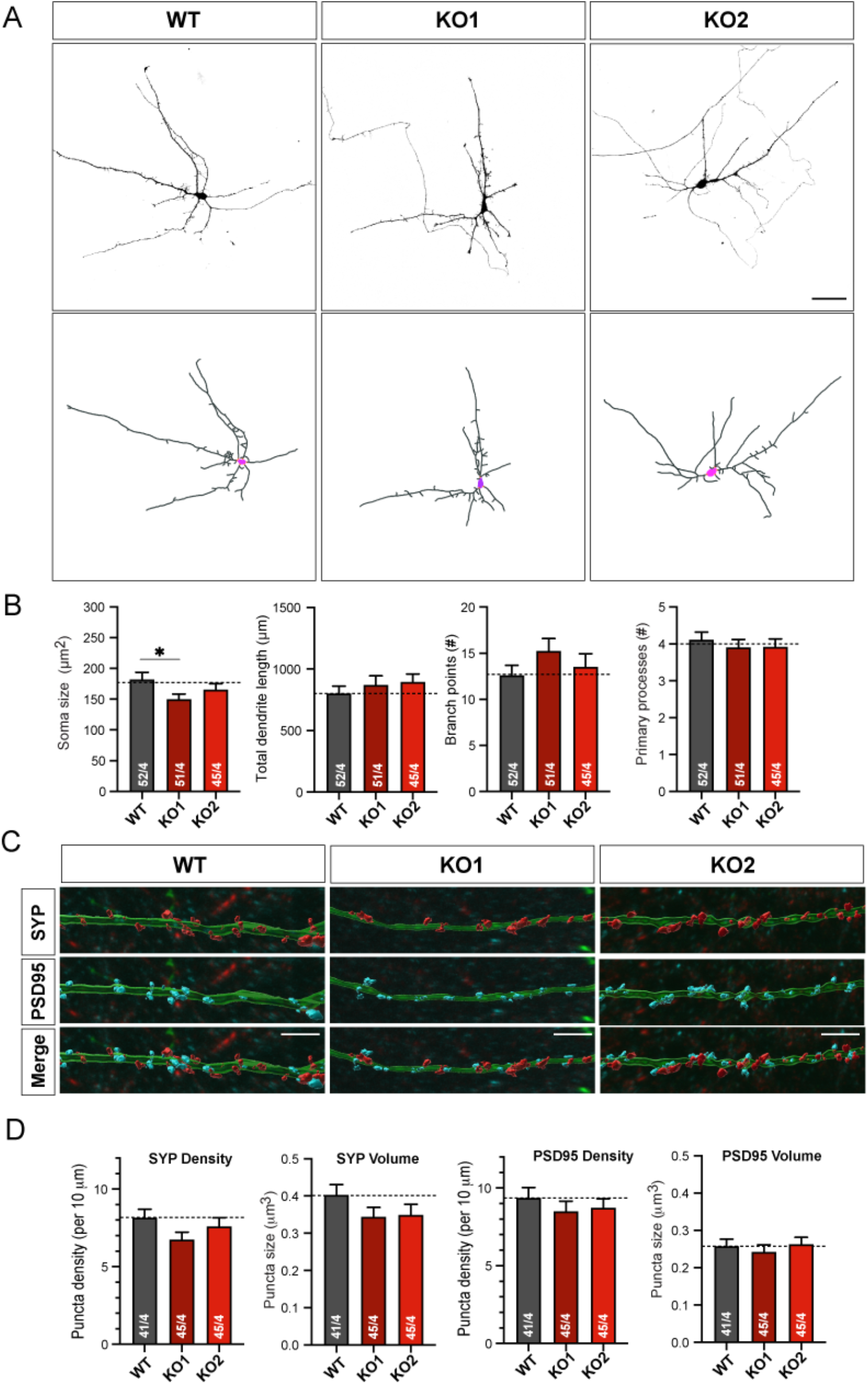
Dendritic arborization and synaptic puncta formation are unchanged in mature day 28 *CASK* KO iNs. (A) Representative images (top) and binary mask (bottom) of iN cultures that were sparsely transfected with SYN-EGFP at day 14 and grown to maturity (day 28). (B) Quantification summary graphs of soma size, total dendritic length, number of branch points, and number of primary processes (scale bar, 50 μm). (C) Binary mask overlaid on top of confocal images showing MAP2+ dendritic segments with pre-synaptic (SYP) and post-synaptic (PSD95) markers (scale bar, 5 μm). Data represents means ± SEM (numbers in bars represent # of cells/# of independent culture replicates performed). Statistical analysis was performed using Student’s t-test comparing WT to each individual KO (*P<0.05; nonsignificant comparisons are not indicated).

### Decreased neuronal network activity in CASK KO mature neurons

To assay neuronal firing at the population level over two different time points (21- and 28-day post neuronal induction), we employed a CMOS-based multi-electrode array system (Maxwell Biosystems) (Müller et al., 2015). In this experimental set-up, we co-cultured WT or *CASK* KO day 3 NGN2 neurons with mouse glia on the recording chips and recorded neuronal activity over two developmental time points, day 21 and day 28 (Fig. 5, Fig. S2-S4). As shown in Figs. 5 and S2-S4, WT neurons showed robust neuronal firing activity as measured by spike firing rate and amplitude at days 21 and 28. The spike amplitude increased over this time period from 60 to 90 μV indicating strengthening of these neuronal connections over development (Fig. 5B, Fig. S3B). In addition to spike activity, WT neurons displayed a consistent level of synchrony as evidenced by the network bursts (Fig. 5C, Fig. S3C), which tend to last longer over time of development in 7 days (0.8 sec in day 21; 1.3 sec in day 28). In *CASK* KO neurons, we observed an overall decrease in the firing rate and amplitude of these neuronal spikes at both days 21 and 28, indicating decreased neuronal firing and strength (Fig. 5, Fig. S3). In addition, the burst frequency, which represents network synchronicity, was significantly lower at both time points and the inter-burst interval was increased (Fig. 5C, Fig. S3C). All other measures of synchrony, such as, mean burst duration, burst peak firing rate, and spikes per burst were not altered, meaning that the composition of bursts is the same (Fig. S4). For both time points, of all the neuronal spikes detected during the recording, there are fewer spikes residing inside these bursts themselves (Fig. 5C, Fig. S3C). Significantly, these firing patterns were observed in two independent *CASK* KO lines and were not due to differences in neuronal plating or active area of the recording chips (Fig. S2). Altogether, these results indicate that although the neuronal morphology and synapse formation are not altered, the strength of these connections as well as their network synchronicity are compromised in *CASK* KO neurons.

**Figure 5.**
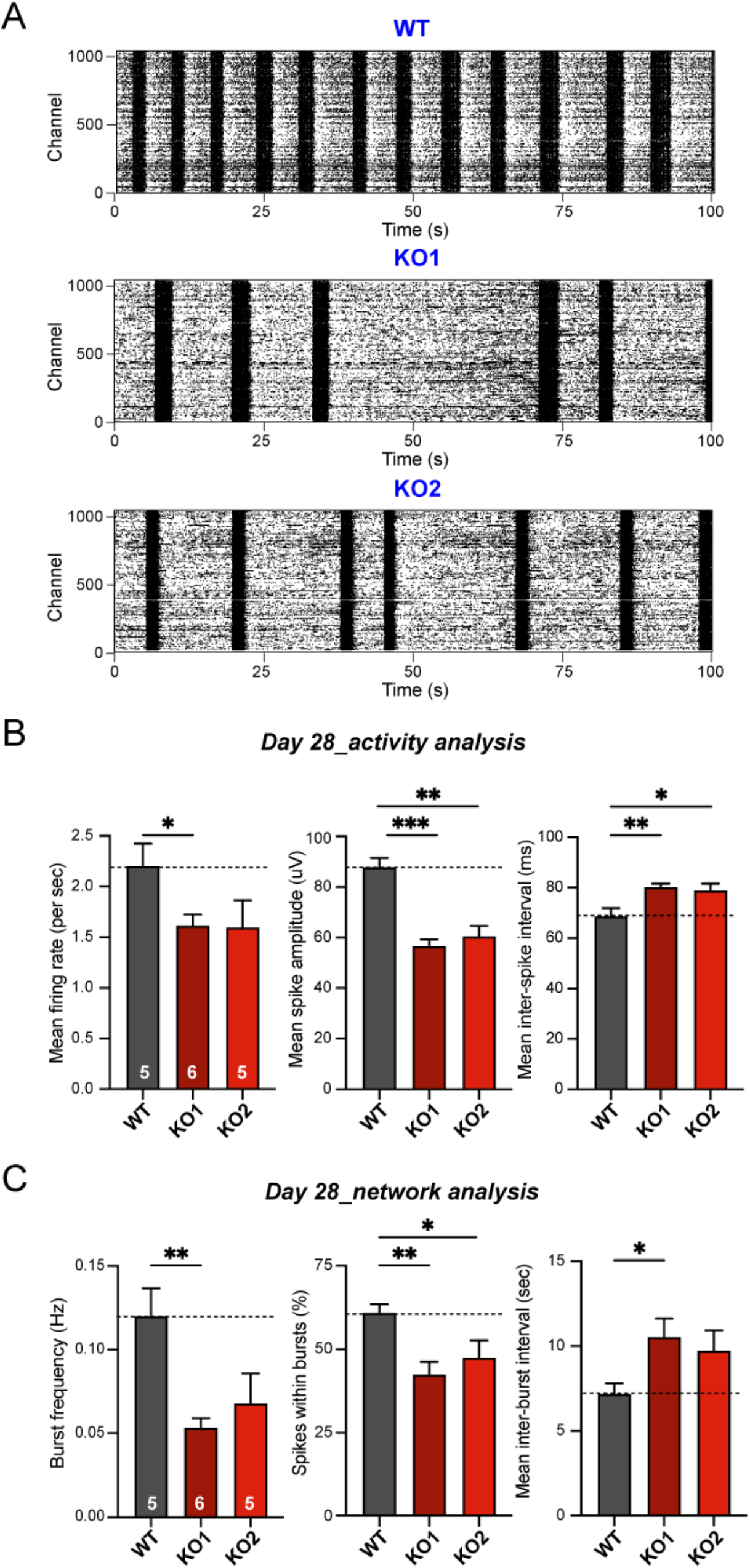
Measurements of synchronous neuronal activity for *CASK* KO mature iNs using high-density microelectrode arrays. (A) Representative raster plots showing decreased network activity for day 28 *CASK* KO iNs as compared to WT. Y-axis represents # of active recording channels and X-axis represents time in seconds (s). (B) Analysis of spike activity revealed a significant decrease in mean firing rate (Hz) and mean spike amplitude (μV), as well as an increase in the mean inter-spike interval (ms) for *CASK* KO neuronal networks across both cell lines. (C) Analysis of network activity indicated a significant decrease in burst frequency (Hz), aligning with an increase in mean inter-burst intervals, and a decrease in the percentage of spikes within bursts in *CASK* KO neurons as compared to WT. Data represents means ± SEM (numbers in bars represent # of independent culture replicates performed). Statistical analysis was performed using Student’s t-test comparing WT to each individual KO (*P<0.05, **P<0.01, ***P<0.001; nonsignificant comparisons are not indicated).

### Defects in synaptic transmission and neuronal excitability in CASK KO mature neurons

We speculated that the observed decreases in MEA spike activity frequency and amplitude could be due to defects in coupling neurotransmitter release machinery and calcium channels at the active zone, a previously documented role for CASK (Maximov and Bezprozvanny, 2002; Olsen et al., 2005). This prompted us to examine properties of synaptic transmission at a single neuronal level using whole-cell patch-clamp electrophysiology (Maximov et al., 2007). Based on the MEA data, we hypothesized that *CASK* LOF would result in a reduction of synaptic release in these cultures. Under voltage clamp configuration, we recorded spontaneous postsynaptic excitatory currents (sEPSCs) in WT and *CASK* KO neurons (KO #1). *CASK* KO neurons showed a profound decrease in the frequency of sEPSCs and not in the amplitude (Fig. 6). Combined with synaptic morphology data, where synapse number was unchanged, this result indicates that there likely is an overall decrease in the probability of neurotransmitter release in *CASK* KO cultures.

**Figure 6.**
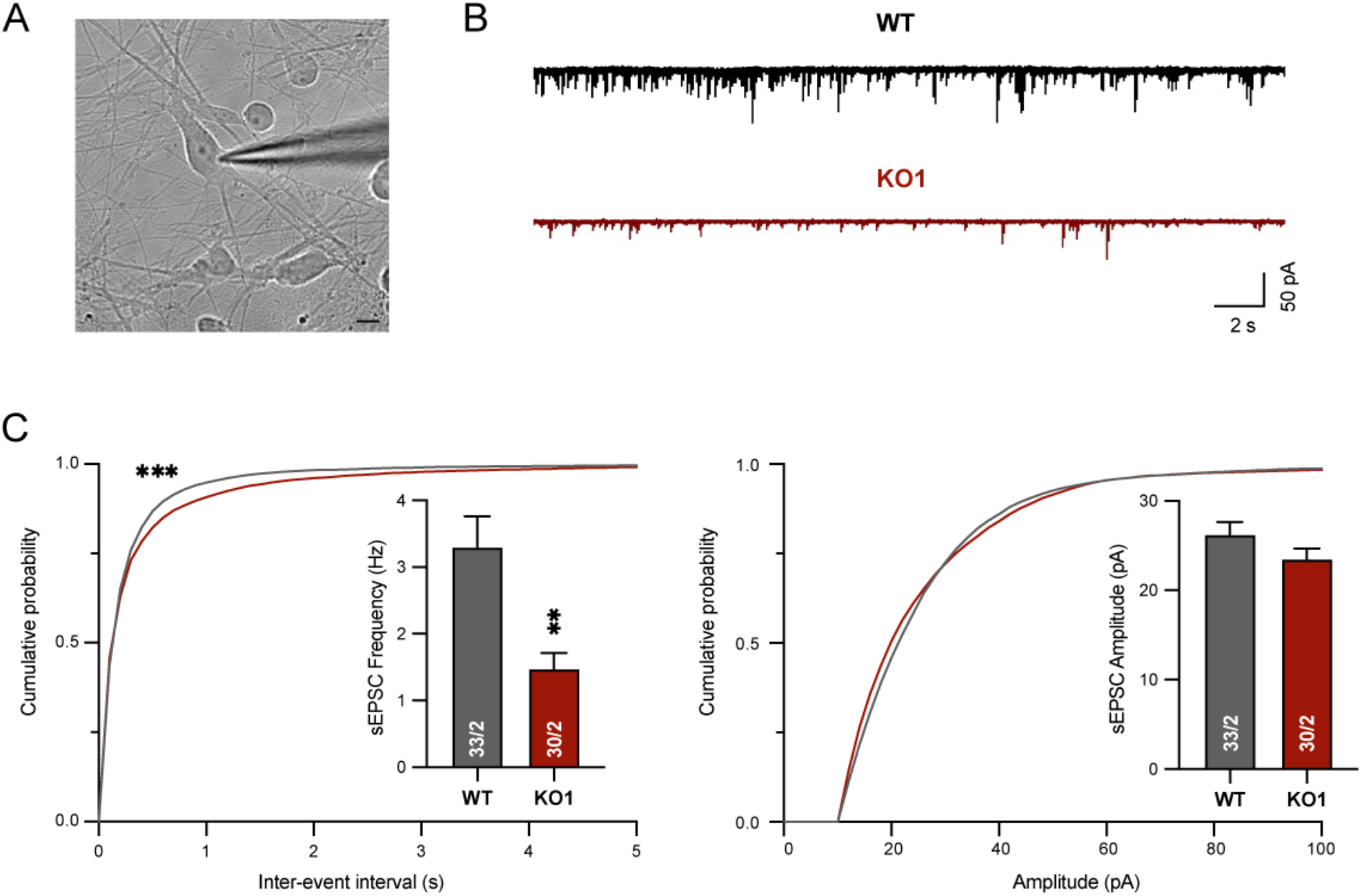
Mature *CASK* KO neurons exhibit a significant decrease in the frequency of spontaneous excitatory synaptic events but no change in amplitude. (A) Representative DIC image of mature iN cells co-cultured with mouse glia with a patch pipette attached to the cell body of the neuron. Electrophysiological recordings in cultured iN cells were performed in the whole-cell patch-clamp configuration (scale bar, 10 μm). (B) Representative traces of sEPSC recordings. (C) Left, cumulative probability plot of interevent intervals and summary graphs of the sEPSC frequency. Right, cumulative probability plot and summary graphs of the sEPSC amplitude. Data represents means ± SEM (numbers in bars represent # of independent culture replicates performed). Statistical analyses were performed by Student’s t-test for the bar graphs, and by Kolmogorov-Smirnov tests for cumulative probability plots, comparing *CASK* KO#1 with control WT neurons (*P<0.05, **P<0.01, ***P<0.001; nonsignificant comparisons are not indicated). All recordings were performed between days 23-28.

We next investigated the intrinsic membrane properties and action potential parameters in neurons generated from the isogenic *CASK* KO and control hES cells under a current clamp configuration. *CASK* KO neurons showed a slight increase in the input resistance, while no statistically significant changes were found in their capacitances (indicative of cell sizes) and their resting membrane potentials (Fig. S5A). The increase in input resistance could be attributed to fewer membrane proteins including ion channels. We then tested their ability to generate action potentials (APs). No obvious changes were found in the *CASK* KO neurons in the major parameters for APs (Fig. S5B), suggesting that LOF of *CASK* likely does not impact the development of voltage-dependent sodium channels as well as potassium channels (two major channels to determine the kinetics of APs). Consistent with the upregulated input resistance of the *CASK* KO neurons, the excitability of these cells is increased because less current injections are needed to depolarize the cell membrane (i.e. *V*_*m*_=*I*_*inj*_\**R*_*in*_) (Fig S5C). The increased excitability could be resulted from the homeostatic changes resulted from reduced synaptic strengths (Fig. 6) and the overall synchrony of firing in the neuronal network (Fig. 5)

In conclusion, based on the results from both whole-cell patch electrophysiology and MEA measurements, there exists a dominant synaptic deficit associated with *CASK* LOF which is driving the decreases in synaptic transmission and network activity. The slight increase in neuronal excitability phenotype cannot compensate for the synaptic driven responses, indicating a potential secondary deficit.

### Differential gene expression networks in CASK KO mature neurons

Given the electrophysiological phenotypes seen in *CASK* KO neurons at both population and single cell level, we investigated the transcriptomic changes associated with *CASK* LOF using bulk RNA-sequencing similar to what had been done in day 7 neurons (Fig. 2). In day 28 neuronal cultures, due to the presence of both human iN cells and mouse glia, we used previously implemented computational methods to bioinformatically separate human iN cells from mouse glial cells, allowing us to identify changes in gene expression specific to human iN cells (Pak et al., 2021). A total of 12 samples were processed and sequenced from four independent cultures collected from WT, KO#1 and KO#2. Using DESeq2 (Love et al., 2014) (FC ≥ 1.2 or FC ≤ 0.8, p ≤ 0.05; minimum of 5 TPM per gene), we identified a total of 1742 DEGs (906 up-regulated, 838 down-regulated) that were commonly dysregulated in both KO#1 and KO#2 compared to WT (Fig. 7A, see SI). A number of DEGs (214 out of 1742) were synaptically localized genes as they mapped to unique SynGO annotated genes, with multiple DEGs assigned to both pre-synaptic and post-synaptic functions (Koopmans et al., 2019) (Fig. 7B). In line with the decrease in the spontaneous synaptic transmission seen in *CASK* KO neurons (Fig. 6), multiple genes regulating synaptic vesicle release were identified as significant DEGs (*VAMP2/SYB2*, *STX1B*, *SYT2*, *RIMBP2*, *RAB3A*, *CPLX1*) (Fig 7B,C). In addition, postsynaptic cell adhesion molecules, neurotransmitter receptors, and signaling molecules (*GRIA2*, *GRIA3*, *GABRA3*, *GABRA5*, *CBLN2*, *SLITRK3-4*, *LRRTM1-3*, *MAP2K1*, *PAK3*, *STAT3*) as well as the components of the ubiquitin proteasome system (*TRIM3*, *NEDD4*, *UBE2A*) were identified as DEGs (Fig. 7B,C), which shows a combinatorial dysregulation of both pre- and postsynaptic gene regulatory network due to *CASK* LOF. Interestingly, previously identified *CASK* targets, *RELN* and NMDA-receptor type subunit (*GRIN1*; for WT vs. KO#1) (Hsueh et al., 2000; Wang et al., 2004a, 2004b), were found to be differentially regulated, further validating these genes as hits. GSEA using Toppgene (Chen et al., 2009) separately validated an enrichment of biological pathways and cellular components involved in synapse organization, assembly, and synaptic signaling (Fig. 7D). DEGs related to neuronal projection development, cellular morphogenesis, and neuronal apoptosis were also enriched (Fig. 7D), despite the normal neuronal and synaptic morphology seen in the *CASK* KO neurons (Fig. 4).

**Figure 7.**
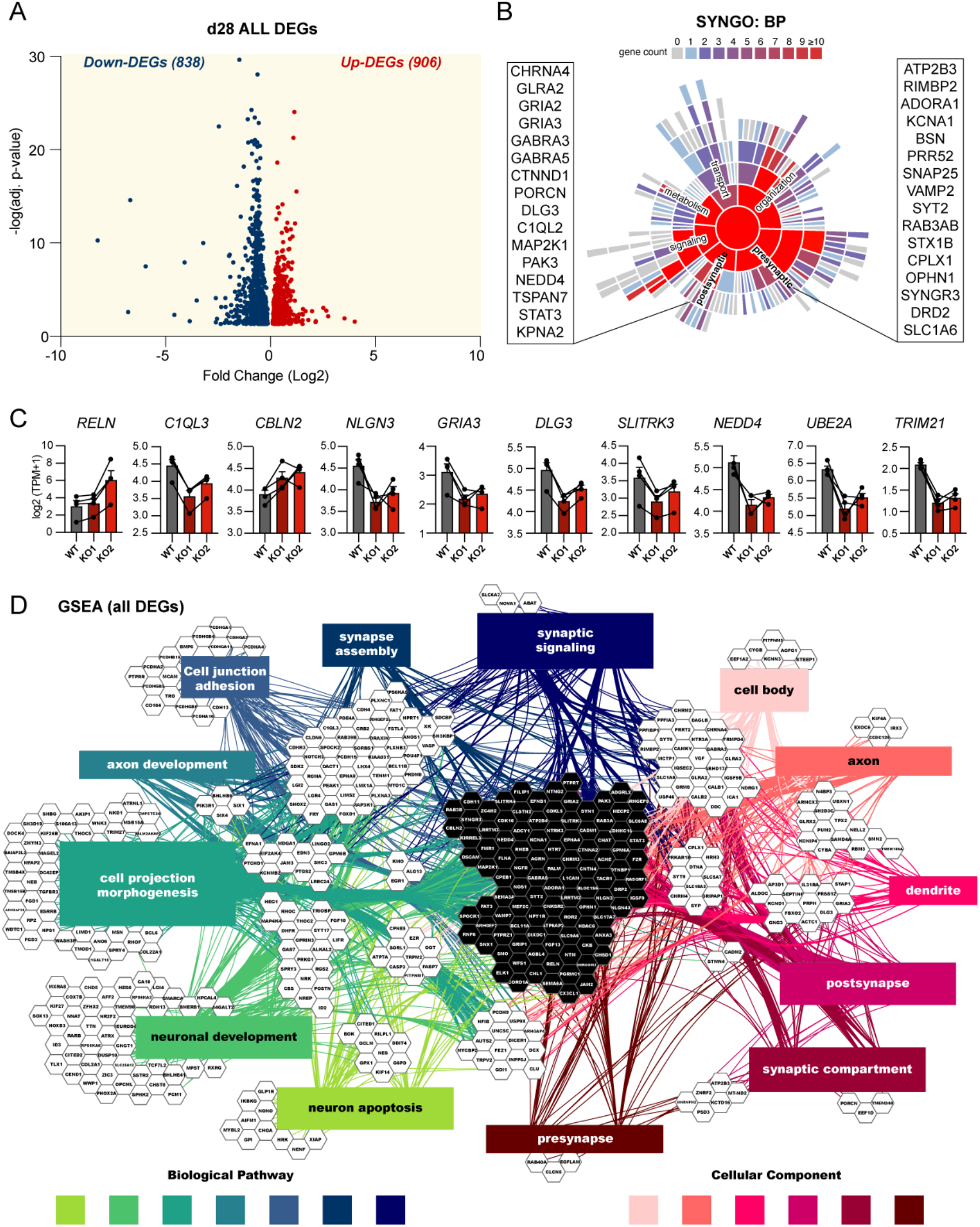
Differential gene expression analysis in d28 *CASK* KO mature iNs using bulk RNA-sequencing. (A) Volcano plot showing −log (adjusted p-value) and log2 (fold change) for overlapping DEGs common in *CASK* KO cell lines compared to WT which met the cutoff for FC ≥ 1.2 or FC ≤ 0.8, p ≤ 0.05 with a minimum of 5 TPM per gene). Paired DEG analysis performed by DESeq2 pipeline (see SI for details). 905 Up-regulated DEGs (red) and 838 Down-regulated DEGs (blue) were chosen for gene set enrichment analysis using ToppCluster and SynGO. All DEGs except *TMSB4X* (log2 FC of −0.58, adj. p-value of 0.0003) was included in the Volcano plot. (B) SynGO-based synaptic gene enrichment analysis identified 214 unique SynGO annotated genes out of 1742 total DEGs. 156 genes have a Biological Process annotation and 189 genes have a Cellular Component annotation. 10 Cellular Component terms are significantly enriched at 1% FDR (testing terms with at least three matching input genes), 5 for Biological Processes. Selected genes pertaining to presynapse and postsynapse are highlighted on either side. (C) log 2(TPM+1) across culture replicates are shown for a selected number of representative DEGs. (D) Gene set enrichment analysis (GSEA; ToppCluster) of overlapping DEGs (FC ≥ 1.5 or FC ≤ 0.5, p ≤ 0.05 with a minimum of 5 TPM per gene) in *CASK* KO#1 and KO#2 compared to WT (Bonferroni p<0.05) identified 7 major biological pathways (BP; synaptic signaling p<4.56E-08; synapse assembly p<1.86E-08; cell junction adhesion p<5.56E-07; axon development p<7.43E-11; cell projection morphogenesis p<1.88E-14; neuronal development p<4.52E-12; neuronal apoptosis p<2.81E-07) and 6 cellular components (CC; presynapse p<2.27E-06; synaptic compartment p<9.18E-15; postsynapse p<2.33E-07; dendrite p<1.35E-08; axon p<2.54E-10; neuronal cell body p<1.69E-07) affected by *CASK* LOF.

## Discussion

Herein, we investigated both the developmental and synaptic roles of *CASK* using human induced neuronal KO models. In an early neuronal growth phase (day 7), we observed an increase in the overall gene expression profiles related to neurite outgrowth, projection morphogenesis, and cell adhesion from transcriptomic analysis and confirmed these changes in morphology as evidenced by increases in neurite length and the number of branch points (Fig. 3). Upon co-culturing these human induced neurons with mouse glia, a culturing scheme which promotes synaptogenesis (Pak et al., 2015; Zhang et al., 2013), we investigated the effects of *CASK* KO on synaptic transmission. As indicated by its previously established protein-protein interactions with neurotransmitter release machinery, voltage-gated calcium channels, and cell adhesion molecule neurexin at presynaptic terminals (Butz et al., 1998; Hata et al., 1996; Maximov and Bezprozvanny, 2002; Mukherjee et al., 2008), *CASK* KO neurons showed a prominent decrease in the frequency of spontaneous EPSCs without changes in neuronal morphology and synapse number (Fig. 4, 6). This result indicated that the regulation of neurotransmitter release is comprised in *CASK* KO cells. In particular, the synaptic deficit seen by whole-cell patch-clamp electrophysiology experiments were supported by synchronous firing activity deficits measured by MEAs (Fig. 5, Fig. S3-4). Indeed, in *CASK* KO cultures, there was an overall decrease in the neuronal spike activity (frequency and amplitude) as well as a decrease in the burst frequency and the percentage of spikes within bursts, indicating that synaptic deficits are driving changes in the network level synchronicity. Although we observed minor changes in neuronal excitability and input resistance, indicating dampened channel activities at the membrane compared to WT (Fig. S5), these parameters are most likely secondary, as these changes cannot compensate for the synaptically driven responses seen at both population and single cell levels (Fig. 5, 6, Fig. S3-4). Interestingly, this phenotype is in line with our previous study in heterozygous *NRXN1* cKO neurons, where a similar decrease in functional connections without changes in morphometric parameters was observed (Pak et al., 2015, 2021), suggesting that functionally, *CASK* LOF mirrors the presynaptic defects driven by *NRXN1* heterozygosity.

In agreement with electrophysiological phenotypes, bulk RNAseq analysis of *CASK* LOF revealed an enrichment of biological functions related to synaptic assembly, synaptic signaling, and cellular components associated with pre- and post-synaptic compartments (Fig. 8). Previously, in rodent models, *Cask* has been shown to regulate transcriptional levels of *Reln* and NMDAR subunit *NR2b* (Hsueh et al., 2000; Mori et al., 2019; Wang et al., 2004a, 2004b), suggesting its role as a transcriptional regulator for its target genes which are important for neurodevelopment. In our transcriptomic data, we also detected an upregulation of *RELN* (~2 fold increase) in both *CASK* KO#1 and KO#2 compared to WT in day 28 neurons (Fig. 8). Additionally, *GRIN1*, which encodes the NMDAR type subunit 1, was upregulated (~1.3 fold increase) in *CASK* KO#1 compared to day 28 WT neurons. The reproducibility of these *CASK* targets in human neurons is encouraging, and our transcriptomic data provides a developmental timing-specific (day 7 and 28) road map of direct and indirect *CASK* targets at the transcriptional level.

Overall, these findings paint a picture of developmental specific functions of *CASK* in human neurons, in agreement with previous findings showing developmentally regulated *Cask* expression patterns and subcellular localization in the brain (Hsueh and Sheng, 1999; Hsueh et al., 2000). In the juvenile brain, *Cask* localizes to the axons to regulate neuronal outgrowth and migration. In the adult brain, *Cask* localizes to synapses to regulate neurotransmitter release and signaling. Our human induced neuronal model at immature and mature time points recapitulated these dual functions of *CASK* by showing developmental timing-specific phenotypes associated with *CASK* LOF. As such, major advantages to this cellular model lie in the ability to 1) assess the functional roles of *CASK* mutations that are relevant to human disorders, 2) bypass organismal lethality seen in model systems due to *CASK* LOF, and 3) examine cell type-specific functions of CASK achieved through neuronal differentiation schemes.

One limitation of the current work is the lack of inhibitory input in the neuronal culture model being studied here. As Ngn2-iN cells are purely excitatory (Zhang et al., 2013), having mixed cultures of both excitatory and inhibitory neurons (Yang et al., 2017) could allow for investigating excitatory-inhibitory (E-I) balance, an important underlying mechanism for ASDs and intellectual disability (Rubenstein and Merzenich, 2003). In two different mouse models, E-I balance defects have been documented in *Cask* mutants, showing an increase in spontaneous miniature excitatory postsynaptic currents and a decrease in spontaneous miniature inhibitory postsynaptic currents (Atasoy et al., 2007; Mori et al., 2019). Mechanistically, the observed E-I imbalance is attributed in part to NMDARs, as re-expression of *NR2b* restores the E-I balance (Mori et al., 2019). Interestingly, a decrease in inhibitory synaptic puncta density was seen in iPSC-neurons derived from ASD *CASK* mutation carriers (Becker et al., 2020). Altogether, these studies show that E-I balance could potentially be disrupted in *CASK* LOF in human cells. From a disease perspective, it will be interesting to test E-I balance in *CASK* mutant context from both isogenic background and diverse disease backgrounds to study the genetic interactions between *CASK* variants and common variants which will likely influence the expressivity of phenotypes mediated by *CASK* deficiency. Nevertheless, using purely excitatory neurons, multiple studies have established the validity of this cellular model for studying the functional consequences of human mutations associated with mental disorders in both isogenic and disease genetic backgrounds and have informed our understanding of phenotypic variation and expressivity (Marro et al., 2019; Pak et al., 2015, 2021; Yi et al., 2016).

In conclusion, our study provides a robust human cellular model for studying the molecular and cellular mechanisms underlying *CASK* deficiency during neurodevelopment. In cortical neurons, *CASK* deficiency elicits developmental timing-specific effects where phenotypes related to neuronal structural integrity and establishment of cortical excitatory neuronal networks are at play. Our results highlight the dynamic nature of *CASK* in human cells, which has multi-functional roles throughout neuronal development and synapse formation. Future studies identifying suppressors of such phenotypes relevant to human patients will provide novel therapeutic strategies for *CASK*-related disorders.

## Experimental procedures

### Pluripotent stem cell culture and gene targeting

H1 human embryonic stem cell (ESC) line (WiCell) was cultured on feeder-free conditions as previously described (Pak et al., 2015, 2021). ESCs were maintained in mTesR medium and passaged using ReLeSR (Stem Cell Technologies). From this wild-type genetic background, CASK knockout lines were generated using CRISPR-mediated gene editing. Briefly, unique sgRNA (5’-CACCGGGACATAGTATTTGAAAGAC-3’) was cloned into lenti-CRISPRv2 construct (Addgene #52961), which expresses Cas9 and puromycin cassette. Lenti-CRISPRv2 plasmid was nucleofected in H1 ESCs using 4D Nucleofector kit (Lonza) and cells were recovered in mTeSR medium. 48 hours of puromycin selection allowed propagation of ESCs that successfully carried lenti-CRISPRv2 construct. Cells were replated for single colony picking and expansion. PCR screen was used to confirm CRISPR-mediated genomic deletions. PCR fragments were validated using Sanger sequencing for positive clones. Two independent KO clones were established (KO#1 and KO#2) which carried 14bp and 10bp deletions respectively which resulted in frameshifts and complete KO. Clones were checked for karyotype and mycoplasma regularly. Neurons were differentiated from working stocks and carried on for no more than ~10 passages.

### Generation of iN Cells from Human ESCs

iN cell differentiation was performed as previously described (Pak et al., 2015, 2021; Zhang et al., 2013). For lentivirus production and primary glia culture, see Supplemental Information for detail. Briefly, on day −1 (d-1), 90-95% confluent ESC cultures were dissociated with Accutase (Innovative Cell Technologies) and plated in 6-well plates at a cell density of 1.0 million cells distributed across 6 wells. Simultaneously, cells along with Ngn2 and rtTA lentiviruses were included in the fresh mTeSR+ plus Y-27632 (Axon Medchem) were used to plate ESCs. On d0, culture medium was replaced with N2/DMEM-F12/NEAA (Thermo Fisher) containing human BDNF (10 ug/L, Peprotech), human NT-3 (10 ug/L, Peprotech), and mouse laminin (0.2 mg/L, Life Technologies). Doxycycline (2 mg/L, Clontech) was also included on d0 to induce TetO gene expression by binding to rtTA and the TetO promoter upstream of the *Ngn2* gene. Doxycycline was included in all subsequent media changes until d10. To select for Ngn2 iN cells, puromycin (1 mg/L, Invivogen) was added to each media change the following two days. Following selection on d3, Ngn2 iNs were co-cultured with mouse glia and plated in 24-well plates (100,000 iN cells/well; 100,000 glial cells/well). Each well contained a Matrigel (BD Biosciences)-coated glass coverslip, and co-cultures were plated in Neurobasal medium (Thermo Fisher) containing B27 supplement, Glutamax (Life Technologies), human BDNF, human NT-3, mouse laminin, and Ara-C (2 uM, Sigma) to inhibit glial proliferation. After co-culture, 50% of the medium was replaced every 2-3 days for each well. iN cells that were cultured up to day 7 followed the same media regimen in the absence of mouse glia. On d10, 50% of medium was replaced by MEM-based maturation medium containing MEM supplemented with 0.5% glucose, 0.02% NaHCO3, 0.1 mg/ml transferrin, 5% FBS, 0.5 mM L-glutamine, 2% B27, and 2 μM AraC. On d13, 50% of medium was supplemented with maturation medium and media change occurred weekly until maturity.

### Immunohistochemistry

Immature d4-transfected iN cell coverslips were harvest on d7, and mature d14-transfected iN cell coverslips were harvested on d28. All coverslips were fixed in 4% paraformaldehyde in PBS+++ (D-PBS containing 0.5 mM CalCl_2_, 1 mM MgCl_2_, and 4% sucrose to minimize coverslip peeling) for 20 min at room temperature and washed three times with PBS+++. For immature d4 iN cells, coverslips were mounted to microscope slides using mounting media Fluoromount DAPI (Southern Biotech). Mature d28 iN cells were further incubated in 0.2% triton X-100 in PBS+++ on a belly dancer for 10 min, and cells were blocked in PBS+++ containing 10% goat serum for 1 hr at room temperature. Primary antibodies used are as follows: rabbit anti-Synaptophysin (1:2000, Abcam ab14692), mouse anti-PSD-95 (1:1000, Invitrogen MAI-046), chicken anti-MAP2 (1:5000, Abcam ab5392), mouse anti-TAU (1:1000, Invitrogen MN1000) and chicken anti-GFP (1:1000, Aves GFP-1010) were added to a mix of 0.1% triton and 5% goat serum and incubated overnight at 4°C. Coverslips were washed three times with PBS+++, and secondary antibodies (1:400) in 5% goat serum were applied for 1 hr. Alexa-546- and Alexa-633-conjugated secondary antibodies (1:1000) were obtained from Invitrogen. Following three final PBS+++ washes, coverslips were mounted to microscope slides with mounting media. For neurite analysis and synaptic puncta measurements, see SI.

### Bulk RNA-Sequencing and Differential gene expression analysis

Four independent culture replicates of immature d7 and mature d28 iN cells were lysed and RNA was extracted using the RNeasy kit (Qiagen). RNA-seq libraries were generated from total RNA using NEBNext Ultra Directional RNA Library Prep Kit for Illumina (NEB). Paired-end sequencing (2 x 75 base-pair reads) was performed using the Illumina Next-seq 500 platform (UMass Amherst Genomics Core), and the resultant fastq files were used for further processing and differential gene expression analysis. For details on DEG generation and GSEA analysis, see SI.

### High-Density Microelectrode Array (HD-MEA) Network Electrophysiology

For preparation of MEA chips, see Supplemental Information. Network activity and burst recordings were performed using Maxwell Biosystems’ CMOS-based single-well MaxOne HD-MEA recording system and accompanying MaxLive software. During each recording, chips were located inside a 37°C, 5% CO2 incubator. An initial activity scan was performed to record the activity by each electrode using the full-scan configuration (20-min recording). Once active electrodes were determined during the activity scan, a 10-min network activity assay was performed including only the electrodes identified as active during the activity scan. For calling signals as “events” during the activity scan, the following thresholds were used: 0.10 Hz (firing rate); 20 μV (amplitude); 200 ms (inter-spike interval). The following network activity thresholding parameters for calling action potentials and network bursts were used: 0.3 sec (smoothing window size); 1.2 (burst detection threshold), 1 sec (minimum peak distance); 0.3 (start-stop threshold). MEA data was exported to Prism (9.3.0) for statistical analysis and graphing.

### Whole-Cell Patch-Clamp Single-Cell Electrophysiology

All electrophysiological recordings for co-cultured iN cells were performed in the whole-cell configuration as previously described and at room temperature (Maximov et al., 2007; Pak et al., 2021). For details, see Supplemental Information.

### Data and code availability

Upon publication, a complete set of bulk RNA sequencing data will be available on NCBI Gene Expression Omnibus (GEO). GSE accession number is in the process of being obtained.

## Acknowledgements

This work was supported by NIMH (R01MH122519 to C.P., R01MH125528 to Z.P.P.), UMass IALS/BMB faculty start up fund (C.P.), Tourette Association of America (Young Investigator Award to C.P.), and NIGMS T32 BTP training program (T32 GM135096 D.M.). We thank Dr. Louise Giam for guidance with CRISPR/Cas9 gene editing, UMass IALS core facilities: Genomics (Dr. Ravi Ranjan) for bulk RNA sequencing assistance and Light microscopy (Dr. Jim Chambers). We also thank members of the Pak lab, Dr. Aldofo Cuadra and Le Wang for fruitful discussion and intellectual support on electrophysiology measurements and Narciso Pavon for assistance with MATLAB code for MEA analysis.

## Author contributions

C.P. generated CRISPR engineered cell lines. D.M. performed all experiments pertaining to iN cell differentiation, morphology analysis, RNA work, and electrophysiology. R.G. performed image analysis and K.J. analyzed RNA-seq data. D.M., Z.P.P., B.A., and C.P. took part in experimental design and data interpretation. D.M. and C.P. wrote the manuscript.

