## Supplementary material for "Loss of Neurodevelopmental Gene *CASK* Disrupts Neural Connectivity in Human Cortical Excitatory Neurons": Supp info

**Supplemental information**

**Supplemental figures and figure legends**

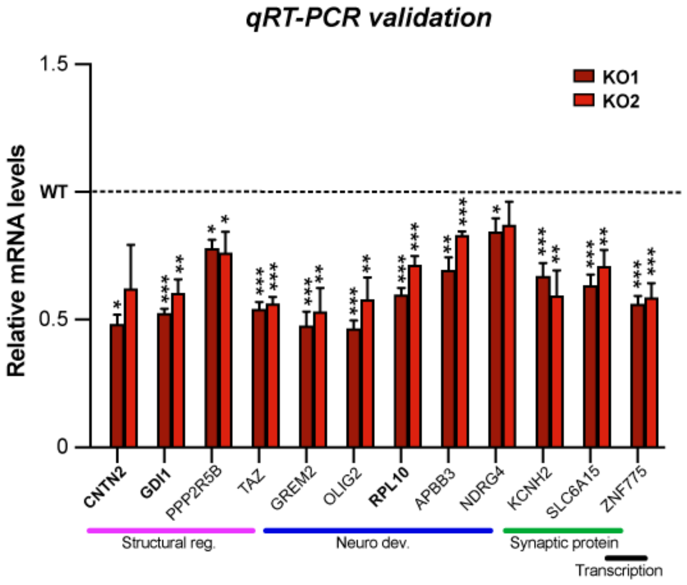


**Figure S1. qRT-PCR validation of down-regulated DEGs.** qRT-PCR validation of specific down-regulated DEGs for both KO#1 and KO#2 relative to WT (set to 1). GAPDH probe was used as a normalization control. Bolded probes indicate genes cross-referenced with SynGO. Four independent replicates per genotype were validated, and each qRT-PCR experiment was performed in triplicate for 12 down-regulated DEGs. qRT-PCR data represents means ± SEM, and statistical analysis was performed using Student's t-test comparing WT to each individual KO (*P<0.05, **P<0.01, **P<0.001; nonsignificant comparisons are not indicated).

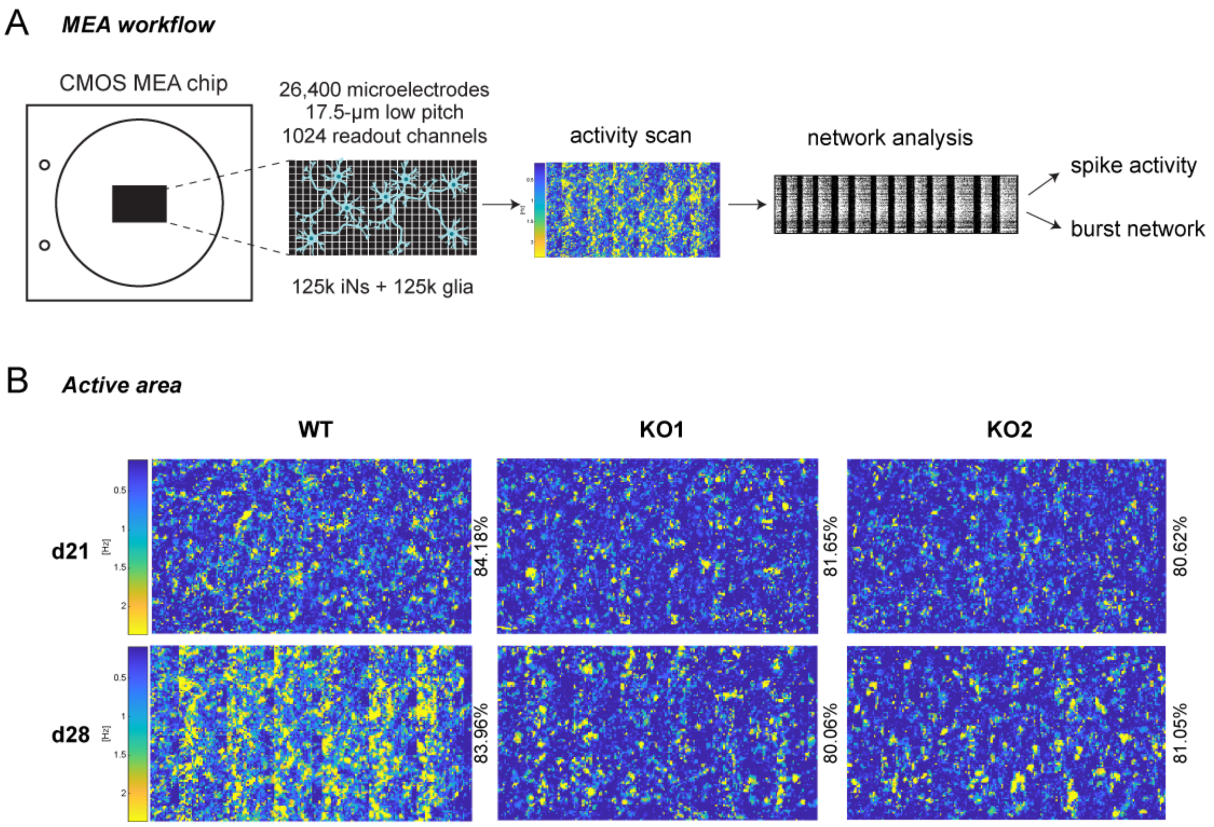


**Figure S2. MEA-based quantification of neuronal network activity.** (A) Schematic showing the workflow of MEA-based neuronal network activity using MaxOne system (Maxwell Biosystems). CMOS-based MEAs chips were plated with 125,000 iN cells + 125,000 mouse glial cells. Each chip was assessed for neural activity (activity scan) and the active areas were then chosen by the software to move forward with network analysis. Each network analysis was subdivided into 1) 'spike activity' to measure mean firing rate (per sec), mean spike amplitude (µV), and mean inter-spike intervals (ms) and 2) 'burst network' to measure burst frequency (Hz), spikes within bursts (%), and mean inter-burst intervals (sec) shown on Fig. 5 and S3. (B) Representative active area maps visualized by activity frequency heatmaps of the entire chip for day 21 and day 28 cultures across genotypes (WT, KO#1, and KO#2). % active area for each chip are shown.

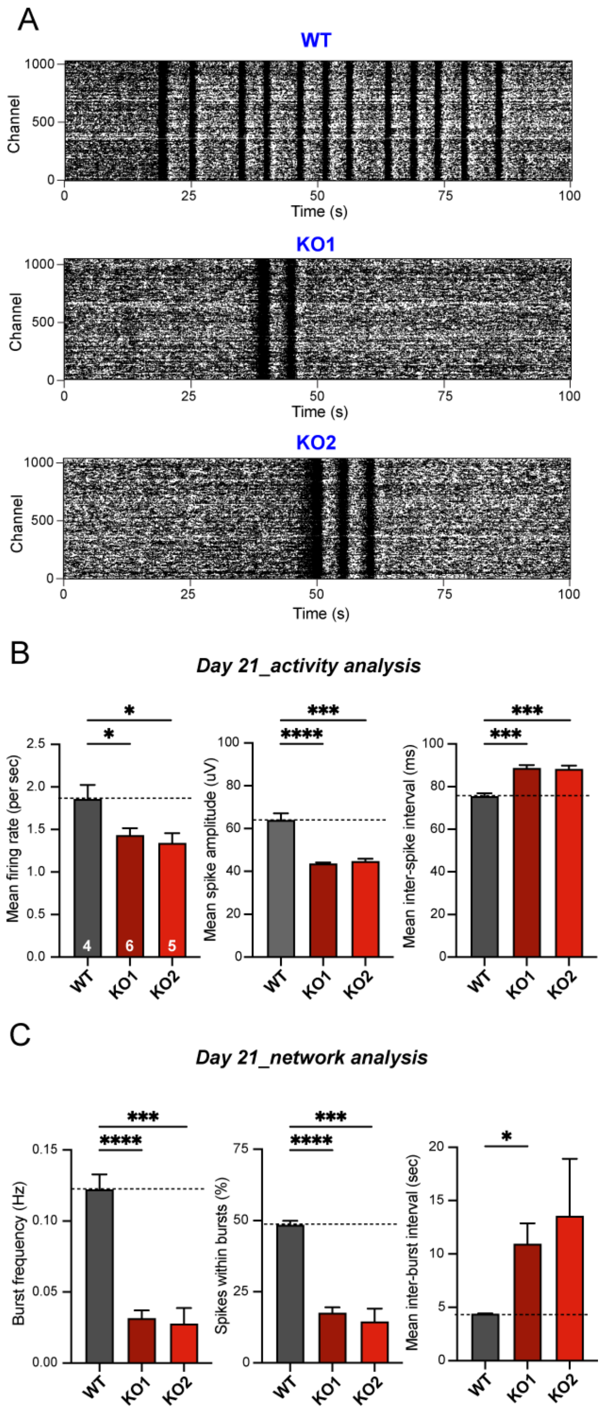


**Figure S3. Measurements of synchronous neuronal activity for *CASK* KO mature iNs using high-density microelectrode arrays**. (A) Representative raster plots showing decreased network activity for day 21 *CASK* KO iNs as compared to WT. Y-axis represents # of active recording channels and X-axis represents time in seconds (s). (B) Analysis of spike activity revealed a significant decrease in mean firing rate (Hz) and mean spike amplitude (μV), as well as an increase in the mean inter-spike interval (ms) for *CASK* KO neuronal networks across both cell lines. (C) Analysis of network activity indicated a significant decrease in burst frequency (Hz), aligning with an increase in mean inter-burst intervals, and a decrease in the percentage of spikes within bursts in *CASK* KO neurons as compared to WT. Data represents means

± SEM (numbers in bars represent # of independent culture replicates performed). Statistical analysis was performed using Student's t-test comparing WT to each individual KO (*P<0.05, **P<0.01, ***P<0.001; nonsignificant comparisons are not indicated).

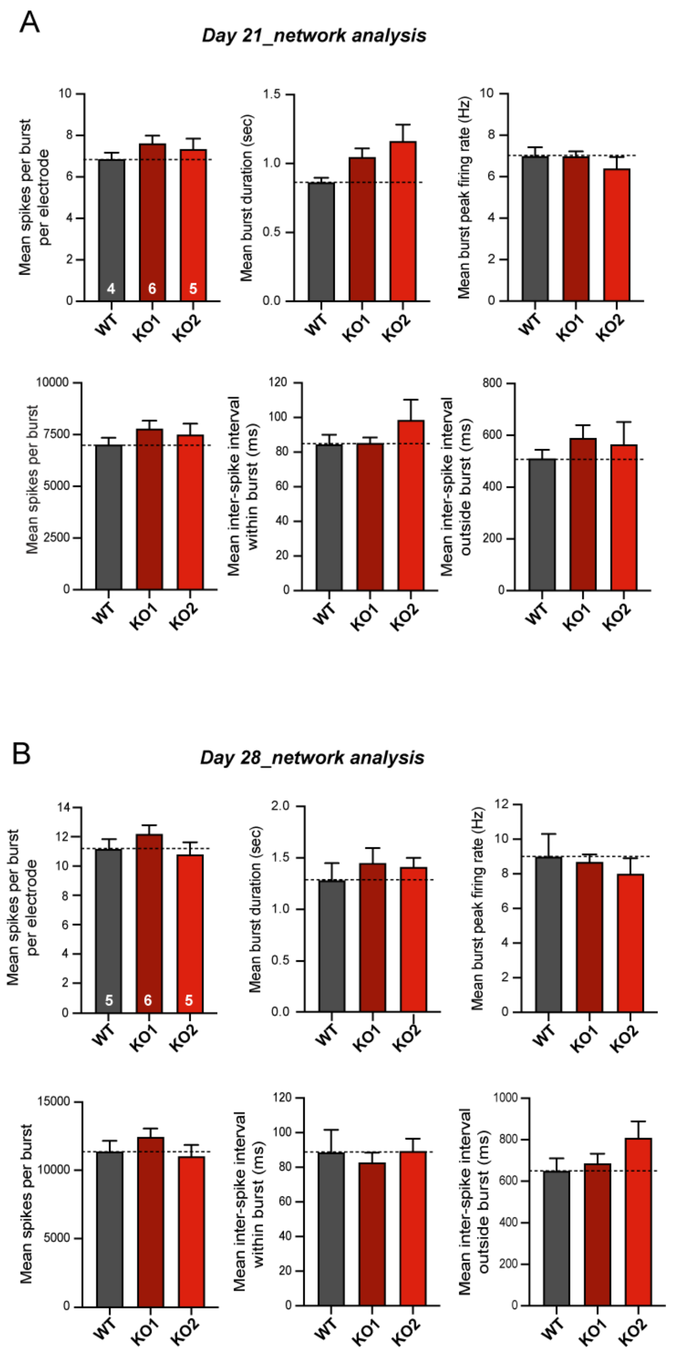


**Figure S4. Quantification of additional parameters of neuronal spikes and synchronous network activities in *CASK* KO mature iNs at day 21 and day 28.** Additional analysis parameters of synchronous network burst activities are shown as means spikes per burst per electrode, mean burst duration (sec), mean burst peak firing rate (Hz), mean spikes per burst, mean inter-spike interval within burst (ms), and mean inter-spike interval outside burst (ms) in day 21 neurons (A) and day 28 neurons (B). There was no significant difference across WT, KO#1 and KO#2 lines. Data represents means ± SEM (numbers in bars represent # of independent culture replicates performed). Statistical analysis was performed using Student's t-test comparing WT to each individual KO (nonsignificant comparisons are not indicated).

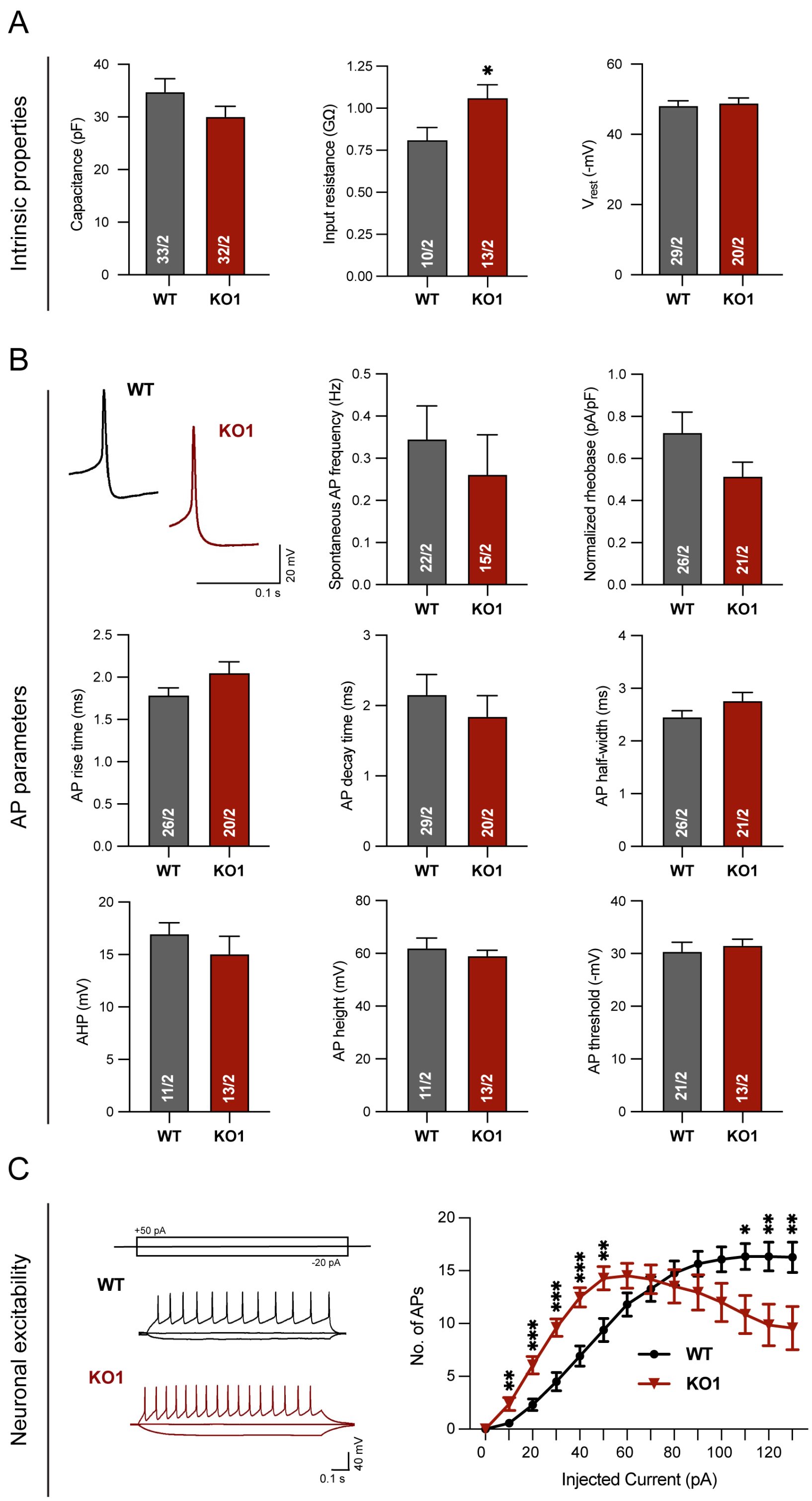


**Figure S5. Measurements of intrinsic electrical properties using whole cell patch clamp electrophysiology.** (A) Intrinsic properties: *CASK* KO#1 neurons show no changes in cell size (capacitance) and resting membrane potential ($V_{rest}$) but show an increase in input resistance compared to WT neurons indicative of immaturity and/or decreased channel activities. (B) Neuronal excitability: *CASK*

KO#1 neurons show an increase in neuronal excitability compared to WT neurons. Left, representative traces of action potentials induced by step current injections, illustration of experimental protocol is shown on top; right, intensity-frequency plots showing the number of evoked action potentials as a function of step current injections. (C) Action potential (AP) parameters: CASK KO#1 neurons exhibit no significant changes in action potential properties. Top left, representative current-clamp recording traces of individual APs. Summary graphs of spontaneous AP frequency, normalized rheobase (rheobase divided by capacitance), AP rise time, AP decay time, AP half width, AP after-hyperpolarization potential amplitude (AHP), AP height, and AP threshold. Data represents means ± SEM (numbers in bars represent # of independent culture replicates performed). Statistical analyses were performed by Student's t-test for the bar graphs comparing *CASK* KO#1 with control WT neurons (*P<0.05, **P<0.01, ***P<0.001; nonsignificant comparisons are not indicated). All recordings were performed between days 23-28.

**Supplemental experimental procedures**

Lentivirus Production

Lentiviruses were produced as previously described (Pak et al., 2021). Briefly, lentiviruses were generated using calcium phosphate transfection of HEK293T cells (Chen and Okayama, 1987) in tandem with three lentiviral packaging plasmids: pRSV-REV, pMDLg/ pRRE and vesicular stomatitis virus G protein expression vector. Per every 75 $cm^2$ of culture area, 12 mg of lentiviral vector DNA was combined with 6 mg of each helper plasmid DNA. 24 hours of incubation following the initial transfection, lentiviruses were harvested from each flask, pelleted by centrifugation (49,000 xg for 90 min), resuspended in MEM, aliquoted, and frozen at -80°C.

Primary glia culture

Primary mouse glial cells were collected from the cerebral cortex of the forebrain of newborn wild-type CD1 mice between postnatal days 0 to 2 (P0-2) as previously described (Pak et al., 2021). Brains were dissected on ice under a light microscope and collected in 1X HBSS (recipe). Mouse cortices were digested using papain (Worthington) and EDTA for 30 minutes and triturated to dissociate cells. Glia were plated on T75 flasks in DMEM with 10% FBS (Hyclone, GE life sciences) and 5% Pen/Strep (Gibco). Once glia reached confluency, cells were trypsinized and plated at 1/3 density. Glia splitting was repeated an additional two times to guarantee the removal of any remaining mouse neurons. Glia were subsequently used for co-culturing with iN cells.

Preparation of MEA Chips

The MaxWell Biosystems CMOS-based single-well MaxOne HD-MEA recording system was used for all network electrophysiology experiments according to the manufacturer's protocols (Maxwell Biosystems, Zurich, Switzerland). Prior to cell plating, MEA chips were first sterilized and pre-coated as follows. First, 1 mL of 1% Terg-a-zyme (Alconox) in deionized water was added to cover each recording array to increase the hydrophilicity of the chip surface. After incubating at room temperature for at least 18 hours, the solution was removed, and chips were rinsed with deionized water. MEA chips were sterilized in 70% ethanol for 30 minutes at room temperature in the biosafety cabinet and washed three times with sterile deionized water. To prepare MEA chips for cell plating, a primary coat of sterile 0.07% poly(ethyleneimine) (PEI) (Sigma/Merck) was applied to each chip and incubated in a 37°C, 5% CO2 incubator for at least one hour. A secondary coat of laminin (0.02mg/mL; Sigma/Merck) was added, and chips incubated at 37°C, 5% CO2 for at least one hour to ensure subsequent cell adhesion to the electrode array. 50 µL of d3 co-cultured cells (150,000 iNs:150,000 glia) suspended in B27-supplemented neurobasal medium were plated on top of the electrode array, and chips were left to incubate at 37°C, 5% CO2 for one hour to allow cells to adhere to the chips. A supplemental 700 µL of B27 medium was added to each MEA well, and chips incubated overnight at 37°C, 5% CO2. Media was changed according to the established Ngn2 culture protocol throughout the experiment, with media change days falling one day prior to each recording.

Quantitative RT-PCR

Total RNA was lysed from cells and extracted as previously described (Pak et al., 2015, 2021). Purified RNA was further treated with RNase free-DNase (Qiagen) to remove genomic DNA contamination and eluted in 40-60 µL per sample. Purified total RNA (~40 µg per reaction) was reverse-transcribed and PCR amplified using Taqman Fast Virus 1-Step Master Mix (Applied Biosystems) and primetime assays for specific genes (IDT). mRNA levels were quantified by real-time PCR assay using the Quantstudio 3 System and RQ analysis software (Thermo Fisher Scientific). qRT-PCR was performed for each of four independent culture replicates for both d7 and d28 neurons. Within each d7 plate, the housekeeping gene *GAPDH* was included as an internal control. Since mouse glia co-cultured with d28 iNs also express *GAPDH*, the neuron-specific *MAP2* gene acted as the internal control for the mature timepoint.

Primer sequences for qRT-PCR
Primetime (IDT) sequences are as follows (probe/forward primer/reverse primer):

**APBB3:**
TGGAGAGTTCACTGGACCGGAGT; CCACCATACCAGGACAGAGA; GGAACTAGGAGATGCAGAGGA

**APLP2:**
AAGAAGGAATGGGAAGAGGCAGAGC; ATGGCTTGGAAGTGCTGAA; CGCAACCGAATGGACAGG

**ARHGAP32:**
AAGAACGTAATGAGCTTGCCGTGC; CACAACCAAACACCCTCTCT; GTGACCAACTCAGTGCCA

**CAMKK1:**
CTGGGCCTCGAGTACTTGCACTG; GATGGCTTGATGTCCCTGT; CAAGCTCGCCTCTACCTG

**CNTN2:**
TTCCAGTAGCGGATCTCATACCCCA; TGCTGTCCTCACTCGGT; ATCCTCAGAGATGAACGTGAC

**CXCL12:**
CGTGCTGGTCCTCGTGCTGA; CATCTGTAGCTCAGGCTGAC; CATGAACGCCAAGGTCGT

**FEZ1:**
CTGCCCTACCTAACCTTGCCCTTT; CTGTGATGGAATGACTCTTGCT; GTGCCTACTTTGCTAACGGA

**GAPDH**:
CAGCAAGAGCACAAGAGGAAGAGAGA; AGGGTGGTGGACCTCAT; TGAGTGTGGCAGGGACT

**GDI1**:
CACAGACATGATGCCCGACAGGAT; GGTGTGATGGAGGAGCTCT; CATGGACGAGGAATACGATGT

**GREM2**:
ACCTTCACCAGCACCGCCA; CTGCTGCCGTCCTTGTAAG; GCTTCCATCACGTCATTGC

**ID1**:
AGAATCTCCACCTTGCTCACCTTGC; CCTGATGTAGTCGATGACGTG;
GCTGTTACTCACGCCTCAAG

**KCNH2**:
AGGACCTGGGTGACCTTCTCAGT; GTGCCTGCAGCTTGTACT; CAGTGACCGTGAGATCATAGC

**KIRREL3**:
CCGACTTCCAGACCATCTACAACTGC; TCCCGACTTCATTTCCGAAC; TGACCATCAGCAACATCGTG

**LRP5**:
ACCTCTCTGAGCCAAGGCCAAAA; GCTGTAGATGTCGATGCTGAG; AGAACATCAAGCGAGCCAA

**LRRC4B**:
CGTCCGGATCACCTGGATGCC; GTTCTTGCTCAGCTGCAGA; CGGTACCTGAACCTGCAAG

**MAP2**:
TCGCAGAGCAGGGAAGAGTGGTA; CAGGAGACAGAGATGAGAATTCC;
CAGGAGTGATGGCAGTAGAC

**NDRG4**:
TATCCCCAACCACCAGCATCACG; CCAGAGTCTGCCATCTTCAG; GGTGCCCAATGCCAAGA

**NECTIN1**:
TACCACTGGACCACGCTAAATGGC; CCTTGAAGAAGAGGGTTCTGTT; CCTGCAAAGCTGATGCTAAC

**NRK**:
CCAACAGCAAATTGACTCCCCACAAAG; GCCTTTCCACTTCTGTCCAT; GAGTTCACTTCTGAGATCTGCT

**OLIG2**:
AGAAGCAAATGACAGAGCCGGAGC; CTTCAAGTCATCCTCGTCCAG; TGTTGATCTTGAGACGCAGC

**PCDHGA10**:
ACTTCTCCATTGGCACCTTCGTCC; ACTTCAACAATTGCGTGTCAG; GTGAGTGTTCCTGAGAATTTGC

**PCDHGB4**:
CCTCCAGTTCCAGCATCACCCT; GCGTTGTCATTTACATCACCAA; CGAGCAGAATCCAGAGTACA

**PCDH8**:
CTTTACCGCTGTCTTTGCCGCTG; GAATCGCTGTCGTTGAAATCAC; CACTCCTTCAACACCATTTCTG

**PLXNB2**:
TCACGGTGTCCAGGTTCTTGCC; CCAGATTCCTGCATGGTCA; CCACGAGACAGATGTGAACTTC

**PPP2R5B**:
ACCACATCTTCCTCCGGTTCATCTATGA; GATCTCCAGCAGCTCAGC; GCCTACATCCGCAAACAGT

**PTPRT**:
TTGCGACTCAAGGTCCGATGCA; AGTTCTCCTGCCAGATCATTC; CACTCTGACTACATCAATGCCA

**ROR2**:
CCTGAGCCGTTATAGCACTGATGGT; CTTGCCGTTCCTCTGTAATCC; GACGCTGCCAACTGCAT

**RPL10**:
CGAGACTTTGGGTACGGCTTGTTCT; TGTACTTATTGGCACAAATTCGG; CACTGAAGATCCTGGTGTCG

**SGK1**:
CATGCCAACATCCTGACCAAGCC; AGTCGTTCAGACCCATCCT; GTGAAGTGAGAGAGCCATGT

**SHANK1**:
ACTTCTTCAACCCCGTCTTCGTGTG; CGAGCTGCACATACTCCAG; TCCGATACAAGACCCGAGTT

**SLC6A15**:
TGTGAACAAAGTTCTGCCACCACCT; ATATTCAGTGCTTCCCTGTACC; TCTCAGTCTTTTCAGCAACCC

**SNCAIP**:
TCCAACGAGATGCCCTGTTCTTGC; CTGTTGCCATCCTGGTCTAC; AGCCATTGCAGAACTGAGT

**SNCB**:
CCACTTCCTCTGGCTTCAGATCAGT; TCCTCATAACTCTCCCCTTCT; CACAGGACTGGTGAAGAGG

**SYT3**:
CCCTCCACGATGTCCCTCCAGA; GCTCCCCAAGATCTGCTTT; TCTATGACTTTGACCGCTTCTC

**TAZ**:
AGGTTCCAGATGTGGCGGAGTTTC; TCCTTGGTGAAGCAGATGTC; CCAATCACCAGTCCTGCAT

**TBC1D30**:
CAATTGTGGGAAAGGGAACGGAGC; ACGGTGTAATGTTGTAGGTGT; CTTCTGCAAAGCCATGAACTC

**UBE2G1**:
TCACCTGCTAATGTTGATGCTGCGA; CTCCATTTCTATCTTCCCTCCA; AGTGTCATTTCTATGCTGGCA

**ZNF332**:
AAGATAGCATTCGTAACAAGAAAGACTCAAGACC; CCTGATCCTGACAATTTGACTTC; CTGAACTAGAGACCTTTTGAAACC

**ZNF445**:
CATGCCATTGCACTCCAGCCTTG; GCAGAAGAATCACTTGAACCCA; GTGGACCGATTGTCGCT

**ZNF737**:
CCTGGTCTTCCTTGGTATTGTTGTCTCT; TGCTCCAGACAGGTGATGA; CTGCACAGCGGAATTTATATAGG

**ZNF775**:
CCAGCTCCTGTGCCGTTGC; TTCTCCTGCTTGACCTTCATC; CGTTAACCTTAGCCACAAAGTC

Sparse transfection in iN cells

To quantify changes in neurite outgrowth, Ngn2 iNs were sparsely transfected with SYN-EGFP construct at either day 4 (for immature iN harvest) or day 14 (for mature iN harvest) using calcium phosphate-based transfection as previously described (Pak et al., 2015, 2021). Briefly, per one well of 24-well format, 0.5 ug of GFP plasmid, 2.5-M $CaCl_2$, and molecular-grade water was mixed and added dropwise to 2X HBS (280 mM NaCl, 1.5 mM $Na_2HPO_4$, 50 mM HEPES) on a vortex, and transfection mix incubated at room temperature for 10 minutes. After washing wells three times with MEM+++ (MEM containing 0.5 mM $CaCl_2$, 1 mM $MgCl_2$, and 4% sucrose to minimize coverslip peeling), transfection mix was added dropwise to each well and incubated for 30 min at 37°C. Wells were washed three final times with MEM+++, and transfection efficiency was confirmed 48 hr later.

Image acquisition and quantification of neurite outgrowth and synaptic puncta

All images were taken with a Nikon A1R25 resonant scanning confocal microscope. Neurite outgrowth was captured using a Zeiss Plan-Apochromat 20×/1.40na Oil DIC Objective M27 (get right info) with 20X magnification at 1024x1024 pixel resolution (aspect ratio: 0.099233 µm per pixel). Synapse formation was imaged using a Zeiss Plan-Apochromat 60×/1.40na Oil DIC Objective M27 (get right info), zoom 2X with 60X magnification and a 2X zoom at 1024x1024 pixel resolution (aspect ratio: 0.099233 µm per pixel). Imaris software (Bitplane), specifically the filamenting and surfaces modules, was used to quantify changes in neurite outgrowth (metrics: number of primary branchpoint, number of branches, total neurite length, soma size), synapse formation, and synapse size. Following parameters were used: Surface Detail- .3 um; Background Subtraction: 1.00 um; Seed point Diameter: 0.3 um. Filter- Number of Voxels: >10. Dendrite to puncta filter-Shortest Distance to Surfaces Surfaces-Dendrite: 0.1 um. Dendrites as close to 150 um in length were chosen to calculate puncta density (but dendritic length ranged greatly). Puncta surface parameters were kept the same for all images. All image quantifications were conducted blindly.

Whole-Cell Patch Clamp Single-Cell Electrophysiology

Patch pipettes were prepared using a Sutter Instruments Model P-1000 Flaming/Brown micropipette puller and borosilicate glass capillary tubes (Sutter Instruments). Intracellular (internal) solution was added to each patch pipette using a microfil needle (World Precision Instruments), and pipette resistance ranged between 3.9-14.7 MOhm. A K-gluconate-based internal pipette solution was used in all experiments, containing (in mM): 126 K-gluconate, 4 KCl, 10 HEPES pH 7.2, 4 Mg-ATP, 0.3 $Na_2$-GTP and10 phosphocreatine; 270-290 mosm/l. The bath solution in all experiments contained (in mM): 130 NaCl, 5 KCl, 2 $CaCl_2$, 1 $MgCl_2$, 10 HEPES (check NaOH and pH), 10 glucose; 290-300 mosm/L. Data were digitized at 10 kHz with a 2 kHz low-pass filter using a Multiclamp 700A amplifier (Molecular Devices), and cellular intrinsic properties of iN cells, including capacitance and input resistance, were recorded in voltage-clamp mode with a minimal amount of current introduced to hold membrane potentials around -70 mV (typically 5-30pA). Spontaneous excitatory postsynaptic currents (sEPSCs) were observed for iN cells in voltage-clamp mode with a holding potential of -70mV and analyzed in Easy Electrophysiology (2.3.3) using the template matching search function and a minimum threshold of 5 pA change in current to call a synaptic event. Action potential parameters, including $V_{rest}$, and spontaneous APs were recorded after break-in but prior to any current injection. To elicit action potentials, a series of increasing amounts of currents was applied to the cell in 5-pA increments across 19 independent sweeps (-50pA to +130pA). Each step lasted for 1 sec, and action potential kinetics were analyzed in ClampFit (9.02) for the first triggered AP observed at the lowest state of current injection.

*Bulk RNA-Sequencing and Differential gene expression analysis*

*Bulk RNA-sequencing & Differential Gene Expression Analysis (d7)*

Raw sequencing reads were processed using an in-house RNA-Seq data processing software Dolphin (Yukselen et al., 2020) at University of Massachusetts Medical School. Paired reads fastq files generated from the NextSeq 500 was imported in the Dolphin analysis suite. The fastq files was subjected to quality check using FastQC. The high quality read pairs first were filtered out using ribosomal RNAs with bowtie2. STAR (2.6.0c) alignment tool (Dobin et al., 2013) was used to align the reads to the reference transcriptome. RSEM (v1.2.28) (Li and Dewey, 2011) was used to quantify the expression levels of genes. The reference transcriptome refSeq annotation, UCSC hg19 refGene was used.

For DEG analysis, top-ranking genes between control and knockout genotypes were filtered for log2(TPM+1)≥1, and a student's t-test (two-tailed unequal variance) was performed. DEGs with a cutoff of p-value>0.05 and |log2FC|>1 were chosen for downstream analysis. K-means clustering and visualization were done in Morpheus.

*Bulk RNA-sequencing (d28)*

On average, 34 million reads per sample were collected using the Illumina Next-seq 500 platform (UMass Amherst Genomics Core). To process paired-end reads, we first built a human + mouse reference (Homo_sapiens GRCh38 v96 and Mus_musculus GRCm38 v96) with a concatenated human-mouse index using the index command in Kallisto (v 0.46.0). d28 RNA-sequencing reads from Fastq files were pseudo-aligned to the concatenated reference genome using the Kallisto quant command and transcripts were assigned to proper species reference. For each Kallisto-processed human-mouse sample, expression matrices of raw count values and TPM (transcript per million) were constructed for human-specific transcripts.

*Differential Gene Expression Analysis (d28)*

Prior to DEG analysis, we first built a transcript-gene conversion table using genecode gene annotation (v39) gtf file (https://ftp.ebi.ac.uk/pub/databases/gencode/Gencode_human/release_39/gencode.v39.annotation.gtf.gz). Transcripts were then loaded and transferred to ensembl gene ids using tximport package (v1.16.1). DESeq2 (v1.28.1) was used to perform DEG analysis for raw count values of 12 d28 samples with the design formula as "design = ~ cell_lines + genotypes", where "cell_lines" is the key of paired samples and "genotypes" is the key of genotype annotations. Significance test results from DESeq2, including log2 fold changes and FDR adjusted p values were used for downstream analysis. Ensembl ids were transferred to gene symbols using biomaRt package (v2.44.4) with Ensembl as the host.

Gene set enrichment analysis (GSEA) for both d7 and d28

ToppCluster (Chen et al., 2009), a multiple gene list feature enrichment analyzer, was used for both d7 and d28 DEG datasets to identify significantly enriched or depleted classes of genes, gene families, or biological pathways affected by CASK-KO. Briefly, the differentially expressed gene dataset was input to the ToppCluster software. 0.05 p-value cutoff using the Bonferroni test was applied to isolate significantly affected gene ontology features; and .xgmml cluster network files were generated and imported into Cytoscape (v3.9.1), an open-source software package for integrating and visualizing these complex molecular interaction and biological pathway networks into string-cluster network maps (Shannon et al., 2003). The enrichment of synaptic genes was analyzed using the SynGO database (Koopmans et al., 2019). For validation of DEGs using qRT-PCR, see Supplemental Information.

Statistics

The experimenter was blind to the genotype of all cultures during imaging analysis and electrophysiological recordings and analysis, and genotypes were unblinded after analysis was complete. Data wrangling was performed in Microsoft Excel, and all raw data points were transferred to Prism (9.3.0) for basic statistics, outlier detection, significance tests, and graph generation. To identify outliers from pooled replicates, the ROUT outlier test was used to identify outliers by fitting data with nonlinear regression and using a false discovery rate of Q=1%. An unpaired parametric two-tailed student t test was performed to compare the two genotypes (WT vs. KO#1 or WT vs. KO#2) for statistical significance. An unpaired nonparametric Kolmogorov-Smirnov test was performed to determine significant differences between the cumulative probability distributions of sEPSC inter-event intervals and amplitudes.

For d7 DEG analysis, top-ranking genes between control and knockout genotypes were filtered for log2(TPM+1)≥1, and a student's t-test (two-tailed unequal variance) was performed. For d28 DEG analysis, paired analysis was done in using the generalized linear model in DESeq2 and p values were generated using Wald test. FDR adjusted p values were generated for multiple comparisons.

**Supplemental tables**

Table S1. List of day 7 DEGs with TPM counts, fold change, and statistics.
Table S2. ToppGene guided GSEA analysis for up- and down-regulated DEGs at day 7.
Table S3. List of day 28 DEGs with raw, TPM counts, fold change, and statistics.
Table S4. ToppGene guided GSEA analysis for up- and down-regulated DEGs at day 28.
Table S5. Mapping of day 28 DEGs to STRING network.
Table S6. SynGO annotation of day 28 DEGs.
